# Aging drives senescence in visceral adipose to a greater extent than obesity, instigating multicellular common signaling patterns that impair organ function and drive inflammation

**DOI:** 10.64898/2026.11.12.765538

**Authors:** Mythili Dileepan, Syed S. Ali, Lee Williams, Mac Anderson, Yuyu He, Juan E. Abrahante Lloréns, Samuel Peters, Yinzhao Wang, Carolina Soto-Palma Erin Hudson, Ananya Raval, Fernanda Rodriguez, Elizabeth L. Schmidt, Erich Kummerfeld, David A. Bernlohr, Christina D. Camell, Constantin Aliferis, Jinhua Wang, Andrew C. Nelson, Paul D. Robbins, Laura J. Niedernhofer, Sayeed Ikramuddin

## Abstract

Obesity accelerates key hallmarks of biological aging in visceral adipose tissue (VAT), yet age-associated changes in cellular senescence against a background of sustained elevated adiposity remain poorly understood. To investigate these changes, we examined human omental VAT from younger (18-35 years), middle-aged (36–51 years), and older adults (52-75 years) with comparable BMI (40 ± 5 kg/m²) using a cross-sectional design. We integrated bulk and single-nucleus RNA sequencing (snRNA-seq), Visium HD spatial transcriptomics, and three complementary computational approaches for senescent cell (SnC) identification. VAT from older adults exhibited increased expression of senescence-associated markers, including *CDKN2A* (p16^INK4a^) and *CDKN1A* (p21^CIP1^), as well as enhanced fibro-inflammatory and tissue-remodeling programs. snRNA-seq revealed a non-linear age-associated pattern, with relatively modest differences between younger and middle-aged adults and more pronounced senescence-associated changes in older adults. SnCs were increased primarily in adipose stem/progenitor cells and mesothelium, which also emerged as major intercellular communication hubs. CellChat analyses identified SnCs of multiple lineages converging on shared outgoing signaling programs, implicating inflammatory, extracellular matrix-remodeling, and adipogenic regulatory pathways, including IL-1, IL-6, TGF-β, and WNT signaling. These predicted interactions were accompanied by transcriptional programs involving HIF1A, NF-κB, SMADs, PRRX1, and TP53 in SnCs, as well as consistent upregulation of *KDM6B* a histone demethylase activating gene expression. Collectively, our findings suggest that aging drives the senescence program to a greater extent than obesity, potentiating tissue dysfunction through coordinated intercellular signaling mechanisms.

## Introduction

Aging and obesity are among the fastest-growing global public health challenges. The number of people aged ≥60 years is projected to reach 2.1 billion by 2050, while the number of adults who are overweight (BMI 25-29.9) or obese (BMI >30) is projected to reach 3.8 billion^1,2^. The convergence of old age and obesity increases the burden of metabolic and cardiovascular disease, frailty, disability and other age-related disorders, creating an urgent need to understand how these processes interact.

Adipose tissue is a dynamic endocrine, metabolic, immune and regenerative organ that undergoes substantial remodeling with aging and obesity. Aging is associated with altered depot distribution, impaired adipogenesis, changes in adipocyte and stromal cell function, immune remodeling, vascular alterations and extracellular matrix (ECM) remodeling. Visceral adipose tissue (VAT) is particularly relevant to metabolic health because its dysfunction is strongly associated with insulin resistance, cardiometabolic disease and systemic inflammation. Aging and obesity share several features of adipose dysfunction, including chronic inflammation, impaired adipocyte differentiation, ECM remodeling, vascular dysfunction and cellular senescence. Senescent cells accumulate in adipocyte, progenitor, endothelial and immune cell populations and alter the tissue microenvironment through inflammatory cytokines, chemokines, growth factors and ECM-modifying factors, contributing to impaired adipogenesis, tissue repair and metabolic homeostasis.

An important challenge is distinguishing molecular changes attributable to chronological aging from those associated with adiposity. Many studies compare individuals with different levels of adiposity or examine aging in metabolically healthy individuals with lower BMI. Such designs are valuable for defining obesity-associated or healthy-aging phenotypes but make it difficult to determine whether aging introduces additional molecular and cellular alterations within an established state of adiposity. Recent single-nucleus (sn) RNA-seq studies have begun to define cellular features of human adipose aging. A prospective study of abdominal subcutaneous adipose tissue (SAT) from 10 younger and 10 older adults, balanced for sex and BMI, identified age-associated adipocyte remodeling, increased immune populations, altered vascular and stromal programs and senescence in specific adipocyte populations; however, participants had BMI of approximately 28-29 kg/m² and the study focused on SAT^3^. Other studies identified age-dependent programs in adipose stem/progenitor cells (ASPCs), including pathways related to adipocyte differentiation and age-obesity interactions, again primarily in SAT^4^. Together, these studies establish important features of adipose aging but leave unresolved how aging remodels human VAT in the setting of established elevated adiposity.

This question is particularly relevant to individuals who develop obesity relatives early in life and remain obese for many years while experiencing progressive metabolic and tissue dysfunction. If aging and obesity share major molecular features, advancing age could amplify an established obesity-associated program. Alternatively, aging may introduce additional cellular and signaling programs that emerge within a stable adiposity state. Addressing this distinction requires a study design that constrains adiposity across age groups.

Because human aging unfolds over decades, directly following the same individuals from young adulthood into older adulthood requires an impractically long longitudinal study vulnerable to participant attrition and other changes over time. Cross-sectional studies therefore provide a practical approach to examining age-associated differences by comparing individuals at different stages of adulthood. However, such studies capture population-level differences between age groups rather than within-person longitudinal trajectories. In the present study, we used a cross-sectional design with comparable high adiposity across young, middle-aged and older female adults to examine age-associated molecular and cellular changes while minimizing differences in adiposity between groups. Here, we investigated human VAT across young (average ∼26y) middle-aged (average-45.4y) and older (average-67y) adulthood in individuals with comparably high adiposity (BMI average 40 kg/m²). We integrated bulk and sn-RNA sequencing to identify SnCs, characterize intercellular communication and define signaling mechanisms impacting tissue function with spatial validation of key cellular features.

A major challenge in studying senescence *in vivo* is the absence of a single universal marker that reliably identifies SnCs. p16^INK4a^ (*CDKN2A*) and p21^CIP^^1^ (*CDKN1A*) remain important components of senescence assessment, but their expression is heterogeneous and can occur in cellular states that do not necessarily represent stable senescence^5^. Thus, p16 or p21 expression alone does not establish a SnC state. More broadly, senescence and their senescence-associated secretory phenotype (SASP) are heterogeneous and tissue-dependent. Here we used three independent computational approaches to identify SnCs in VAT of obese adults, which all converged on ASPCs and mesothelial cells being the major contributors. Transcriptomic analysis of these SnC populations revealed common patterns of gene expression rewiring driving inflammation (increased NF-κB signaling), extra-cellular matrix remodeling (increased SMADs and PRRX1 activity), hypoxia (elevated HIF1A activity) and impaired adipogenesis (increased WNT signaling), with evidence of epigenetic regulation via *KDM6B*.

## Methods

### Human tissue collection and ethical approval

Human adipose tissue samples were obtained through the University of Minnesota from patients undergoing bariatric surgery, archived tissue specimens maintained by the University of Minnesota BioNet Tissue Procurement Facility, and deceased organ donors through LifeSource, an Organ Procurement Organization (OPO).

Prospective tissue collection from patients undergoing bariatric surgery was coordinated through the University of Minnesota Surgery Clinical Trials Office (SurgCTO) and BioNet. Tissue samples were collected from patients who provided informed consent for tissue donation and research use. Additional archived human adipose tissue specimens were obtained from the BioNet tissue repository in accordance with the applicable institutional research protocols and consent requirements.

Tissue samples from deceased organ donors were obtained through LifeSource following applicable donor authorization and institutional procedures for research tissue procurement. The collection and use of surgical and archived BioNet tissue samples were approved by the University of Minnesota Institutional Review Board (IRB; Protocol No. [STUDY00021497]). All tissue samples were assigned coded identifiers, and available donor demographic and clinical information, including age, sex, body mass index (BMI), and relevant medical history, was collected in accordance with the approved protocols. All procedures were conducted in compliance with applicable institutional and ethical guidelines for research involving human biological specimens.

### Study design and donor characteristics

Omental visceral adipose tissue (VAT) samples were collected from adult female donors aged 18-75 years with a body mass index (BMI) of 40 ± 5 kg/m². Donors were categorized into three age groups: young adults (YA; 18-35 years), middle-aged adults (MA; 36-51 years), and older adults (OA; 52-75 years). At least 3-6 biological samples were included per age group, depending on the experimental analysis. This study design allowed us to investigate age-associated changes in cellular senescence and intercellular communication in human omental VAT among donors with comparable adiposity.

We analyzed publicly available datasets from Jung *et al.* (2026)^6^, Lazarescu *et al*. (2025)^7^, and Emont *et al.* (2022)^8^ to compare young and older adults with low BMI, as well as young adults with low and high BMI. For the age-related comparison, we included young adults (18-35 years) and older adults (52-75 years) with low BMI.

### Adipose tissue collection, processing, and storage

Fresh human omental visceral adipose tissue (VAT) tissues were transported to the laboratory on ice in cold Dulbecco’s Modified Eagle Medium (DMEM), while frozen specimens were transported on dry ice and stored at −80°C until further analysis.

For histological studies, tissues were fixed in 10% neutral-buffered formalin at 4°C for approximately 36 hours, washed with distilled water, and stored in 70% ethanol until formalin-fixed, paraffin-embedded (FFPE) processing.

When sufficient fresh tissue was available, stromal vascular fraction (SVF) cells were isolated using a gentleMACS Octo Dissociator and the Adipose Tissue Dissociation Kit (Miltenyi Biotec, Cat. No. 130-105-808), following a laboratory-adapted protocol for human adipose tissue. Isolated SVF cells were cryopreserved in medium containing 10% dimethyl sulfoxide (DMSO) using a Mr. Frosty freezing container at −80°C before transfer to liquid nitrogen for long-term storage. Frozen tissues were used for bulk RNA-seq, single-nucleus RNA-seq (snRNA-seq), and other molecular analyses, while tissue sections were used for spatial transcriptomic and histological studies.

### Bulk RNA-seq: RNA isolation, library Preparation, and sequencing

Total RNA was extracted from frozen human omental visceral adipose tissue (VAT) using a modified TRIzol-RNeasy Plus Mini Kit protocol. Briefly, tissue samples were homogenized in TRIzol reagent (Invitrogen, Thermo Fisher Scientific), followed by chloroform-induced phase separation according to the manufacturer’s instructions. The RNA-containing aqueous phase was collected and further purified using the RNeasy Plus Mini Kit (QIAGEN) following a laboratory-adapted protocol. RNA quality and integrity were assessed using an Agilent 2100 Bioanalyzer (Agilent Technologies).

Eight uniquely dual-indexed (UDI) stranded RNA-seq libraries were prepared using Takara SMARTer Pico Mammalian RNA-Seq Kit v2 reagents. The pooled libraries were gel size-selected to obtain approximately 200-bp inserts and sequenced on an Illumina NextSeq 2000 platform using a P2 flow cell with 150-bp paired-end reads (2 × 150 bp). The sequencing run generated up to 400 million reads, with all expected barcodes detected and well represented. All libraries achieved mean sequencing quality scores of Q30 or higher.

### snRNA-seq: Nuclei isolation, library preparation, and sequencing

Nuclei were isolated from frozen human omental visceral adipose tissue (VAT) using a published protocol, Nuclei Isolation from Human Frozen Adipose Tissue for Single-Nuclei RNA Sequencing (version 2; DOI: 10.17504/protocols.io.eq2ly6w5qgx9/v2), adapted from three previously published methods. Isolated nuclei were processed without fixation using the standard 10x Genomics Chromium single-nucleus RNA sequencing (snRNA-seq) workflow. Single-nucleus gene expression libraries were prepared using the Chromium Single Cell 5′ Gene Expression platform (10x Genomics), targeting approximately 20,000 nuclei per sample. Libraries were sequenced on an Illumina NovaSeq platform at a target depth of approximately 50,000 reads per nucleus. Donor metadata are presented in **Supplementary Table 1**.

### Single-nucleus RNA sequencing data processing and analysis

FASTQ files generated from 10x Genomics single-nucleus RNA sequencing (snRNA-seq) were processed using Cell Ranger (v8.0.1)^9^ and aligned to the human GRCh38 reference genome obtained from the 10x Genomics website. The filtered gene expression count matrices generated by Cell Ranger were imported into Seurat (v5)^10^ for downstream analysis. Mitochondrial gene expression was calculated for each nucleus, and nuclei with 500-4,000 detected genes (‘nFeature_RN’) and less than 5% mitochondrial gene expression were retained^10^.

Raw counts were normalized using the ‘NormalizeData ()’ function. For sample integration, 2,000 highly variable genes were identified using ‘FindVariableFeatures ()’ with the variance-stabilizing transformation (VST) method^10^. Integration anchors were identified using a reference-based approach, with one sample designated as the reference, followed by integration using ‘IntegrateData ()’^10^. The integrated assay was scaled using ‘ScaleData ()’, and principal component analysis (PCA) was performed using default parameters^10^. The first 15 principal components (PCs) were used for nearest-neighbor identification, clustering, and uniform manifold approximation and projection (UMAP) visualization^10^. To improve cluster separation and minimize potential batch effects, Harmony^11^ was subsequently applied, and the resulting Harmony components were used to generate the final clusters and UMAP projections.

### Cell-type annotation and subclustering

Major cell populations, including adipocytes, macrophages, adipose stem and progenitor cells (ASPCs), endothelial cells, and mesothelial cells, were individually extracted for subclustering analysis. For each population, PCA, nearest-neighbor identification, clustering, and UMAP projection were repeated using the first 15 components. Clustering resolutions ranged from 0.2 to 0.5, depending on the cell population. Cell clusters and subclusters were annotated based on established marker genes and the human adipose tissue atlas published by ^8^ Emont *et al*., 2022.

### Identification of predicted senescent cell populations

Three complementary computational approaches were used to identify putative SnCs: (1) gene selection based on SEAC-defined categories (2) SEAC reference-based correlation analysis, (3) SenMayo gene signature scoring **(Supplementary Table 2).**

### SEAC-based senescent cell identification

A complementary approach was used to identify nuclei exhibiting coordinated enrichment across four senescence-associated functional programs

1. **S**enescence-associated secretory phenotype (SASP): Positive enrichment of IL1B, IL6, CCL7, CXCL8, CCL2, CXCL2, and CD55.
2. **E**xtracellular matrix remodeling: Positive enrichment of TMEM217, ADAMTS4, VCAM1, PLAUR, PLAU, TNC, and ICAM4.
3. **A**nti-apoptotic signaling: Positive enrichment of SERPINE1, SERPINB2, BCL2L1, and MCL1, alongside negative enrichment of GZMB and RELT.
4. **C**ell-cycle regulation: Positive enrichment of CDKN1A, TP53, GADD45A, and CDKN2A, alongside negative enrichment of CCNB1, CCNB2, and CDK1.

Gene signature scores were calculated using UCell^12^, incorporating the specified positive and negative gene-expression directions. Nuclei exhibiting positive enrichment scores across all four functional categories were classified as putative senescent cells. Specifically, cells were required to exceed a score threshold of 0.1 across all four categories: SASP, extracellular matrix remodeling, anti-apoptotic signaling, and cell-cycle regulation. Thresholds were selected based on score distributions, identifying populations residing in the tails of each plotted distribution.

### SEAC-Reference-based correlation analysis

Cells co-expressing the senescence-associated genes CDKN1A, BCL2A1, MCL1, SERPINE1, ICAM1, PLAUR, PLAU, CCL2, IL6, and CXCL2 and assigned to the G1 phase using Seurat were identified as a reference cell pool. The second cell in the ordered list of this pool was selected as the reference cell. Pearson correlation coefficients were then calculated between the gene expression profile of the reference cell and that of each nucleus. Nuclei exhibiting a Pearson correlation coefficient > 0.55 were classified as putative senescent cells.

### SenMayo gene signature scoring

The published SenMayo gene signature¹³ was used to assess senescence-associated transcriptional activity. Gene signature enrichment scores were calculated for individual nuclei using UCell¹², and nuclei with scores exceeding the 99th percentile were classified as putative senescent cells. The threshold was determined using the R quantile () function with probs = 0.99.

### Differential gene expression analysis

Differential gene expression analysis was performed to identify age-associated transcriptional changes by comparing older adults (OA) with young adults (YA) and middle-aged adults (MA). Analyses were conducted across the combined dataset and separately within major cell populations, including ASPCs, mesothelial cells, adipocytes, macrophages, and endothelial cells. Differentially expressed genes (DEGs) were identified using Seurat’s ‘FindMarkers()’ function with default parameters^10^

To identify transcriptional differences between putative senescent and non-senescent cells, a repeated subsampling strategy was implemented to minimize potential bias arising from differences in cell population sizes. Within each cell type, non-senescent nuclei were randomly sampled to match the number of putative senescent nuclei. Differential gene expression analysis was performed between these equally sized populations, and the procedure was repeated 100 times. Genes meeting the specified adjusted P-value threshold of [insert threshold], following Benjamini-Hochberg correction, were retained for downstream analyses.

### Cell-cell communication analysis

Cell-cell communication was inferred using the ‘CellChat’ R package^14^ to characterize ligand–receptor interactions and signaling pathways across age groups. Analyses were performed using the standard CellChat workflow and default parameters. Predicted ligand-receptor interactions and associated signaling pathways meeting the significance threshold of P < 0.05 were retained.

To investigate senescence-associated intercellular communication, the OA dataset was further analyzed by distinguishing putative senescent and non-senescent cell populations. The standard CellChat workflow was repeated following the procedures described in the CellChat GitHub documentation^14^. Predicted outgoing and incoming signaling interactions were examined to characterize communication between senescent cells and other cell populations.

### Transcription factor activity analysis

Transcription factor (TF) activity was inferred using the ‘decoupleR’ R package^15^. A univariate linear model was applied to estimate transcription factor activity based on gene expression and regulatory interactions. Inferred TF activity scores were evaluated as relative measures of regulatory activity, and transcription factors exhibiting higher activity scores in putative senescent cells were prioritized for downstream interpretation.

### Visium HD spatial transcriptomics and data analysis

Visium HD spatial transcriptomic analysis was performed on eight human omental visceral adipose tissue (VAT) samples from female donors, including four young adults (YA) and four older adults (OA). Formalin-fixed, paraffin-embedded (FFPE) tissue sections (7 µm thick) were mounted on NEXTERION Slide H 3-D hydrogel-coated glass slides (SCHOTT North America, Cat. No. NC0782819) and processed for Visium HD spatial gene expression analysis (10x Genomics). Libraries were sequenced using the AVITI sequencing platform (Element Biosciences). Raw sequencing data were processed using Space Ranger (v4; 10x Genomics) to generate spatial gene expression matrices.

Spatial transcriptomic data were analyzed at a 16-µm bin resolution using the Scanpy Python library. Gene expression counts from each sample were normalized and log-transformed, followed by highly variable feature selection. Samples were integrated using Harmony to minimize batch-associated variation. Leiden clustering was subsequently performed on the integrated data, and cluster-specific marker genes were identified using Scanpy to annotate putative cellular identities of the spatial bins. Cluster annotations were validated by overlaying the spatial distribution of annotated bins onto the corresponding tissue microscopy images.

### COMET multiplex immunofluorescence staining and imaging

Multiplex immunofluorescence staining was performed on formalin-fixed, paraffin-embedded (FFPE) human omental visceral adipose tissue (VAT) sections (5 µm thick) using the COMET spatial imaging platform (Lunaphore). Tissue sections were processed according to the manufacturer’s instructions, with an additional laboratory-adapted blocking step. Briefly, following antigen retrieval and before the first antibody staining cycle, tissue sections were incubated with 10% goat serum for 10 minutes at room temperature (RT) to minimize nonspecific antibody binding. Subsequent multiplex immunofluorescence staining and imaging were performed according to the manufacturer’s standard COMET protocol. The following primary antibodies were used: anti-CDKN2A (Abcam, Cat. No. ab108349, clone EPR1473), anti-CDKN1A (Cell Signaling Technology, Cat. No. 2947S, clone 12D1), anti-KDM6B (R&D Systems, Cat. No. AF7300), anti-CD31 (R&D Systems, Cat. No. MAB11666-100, clone 1094431), anti-cytokeratin (Lunaphore, Cat. No. MR10140, clone AE1/AE3), and anti-γH2AX (Cell Signaling Technology, Cat. No. 203045S, clone D7T2V). The secondary antibodies used were anti-rabbit Alexa Fluor 647 (Lunaphore) and goat anti-mouse IgG (H+L), Alexa Fluor 750, cross-adsorbed (Thermo Fisher Scientific).

### RNA extraction, reverse transcription, and quantitative real-time PCR

Total RNA was extracted from human omental visceral adipose tissue (VAT) using the RNeasy Plus Mini Kit (QIAGEN, Cat. No. 74134) according to the manufacturer’s instructions. Complementary DNA (cDNA) was synthesized from 1 µg of total RNA using the High-Capacity cDNA Reverse Transcription Kit (Applied Biosystems, Thermo Fisher Scientific, Cat. No. 4374967) following the manufacturer’s protocol. Quantitative real-time PCR (qPCR) was performed using TaqMan Gene Expression Assays (Applied Biosystems) on a QuantStudio 5 Real-Time PCR System (Applied Biosystems). Expression levels of the senescence-associated genes *CDKN2A* (p16) and *CDKN1A* (p21), and the inflammatory gene *TNF* (TNF-α), were measured using 5–7 independent biological samples per age group. Relative gene expression was calculated using the 2^−ΔΔCt method, with 18S ribosomal RNA (18S rRNA) as the endogenous reference gene and the young adult (YA) group as the calibrator.

### Statistical analysis

Statistical analyses were performed using GraphPad Prism version 11 (GraphPad Software, Boston, MA, USA) and R (version 4.4). Independent human donors were considered biological replicates, with sample sizes ranging from 3 to 5 donors per age group, depending on the experiment. Data are presented as mean ± standard error of the mean (SEM), unless otherwise indicated. Differences among young adults (YA), middle-aged adults (MA), and older adults (OA) were evaluated using one-way analysis of variance (ANOVA), followed by Tukey’s multiple-comparisons test. Ordinary two-way ANOVA was used to evaluate the effects of age group and cell type, including their interaction, on the proportions of predicted senescent cells. Post hoc multiple-comparisons tests were performed to assess differences between age groups within individual cell types.

For single-nucleus RNA sequencing (snRNA-seq), differential gene expression analysis was performed using Seurat’s ‘FindMarkers ()’ function. P values were adjusted for multiple testing using the Benjamini-Hochberg false discovery rate (FDR) method. Gene Ontology enrichment analysis was performed using clusterProfiler, with FDR correction applied for multiple comparisons. An adjusted P value <0.05 was considered statistically significant for transcriptomic analyses, whereas P <0.05 was considered significant for other statistical comparisons.

## Results

### Aging-related senescence in visceral adipose tissue preferentially occurs in adipose progenitor and mesothelial cell populations

We first determined whether aging is associated with molecular and cellular changes in visceral adipose tissue (VAT) senescence among individuals with comparable levels of obesity. VAT samples were collected from individuals across different age groups with similar BMI (40 ± 5 kg/m²), and transcriptomic analyses was first performed at the bulk RNA level in younger adults (18-35 years; mean age, 26 years; n = 4) and older adults (52-75 years; mean age, 67 years; n = 4). **(Fig. 1a).** Bulk RNA-seq revealed robust upregulation of senescence-associated biomarkers in VAT from older adults compared with younger adults **(Fig. 1b),** together with enrichment of pathways associated with canonical features of cellular senescence **(Fig. 1c and Extended Fig. 1)**. Consistent with these findings, the bulk transcriptome of older obese adults showed a positive correlation with the SenMayo gene panel, whereas that of younger obese adults showed a negative correlation **(Fig. 1d)**. Together, these findings demonstrate an age-associated increase in senescence-related transcriptional programs in VAT in a setting of obesity.

**Figure 1.**
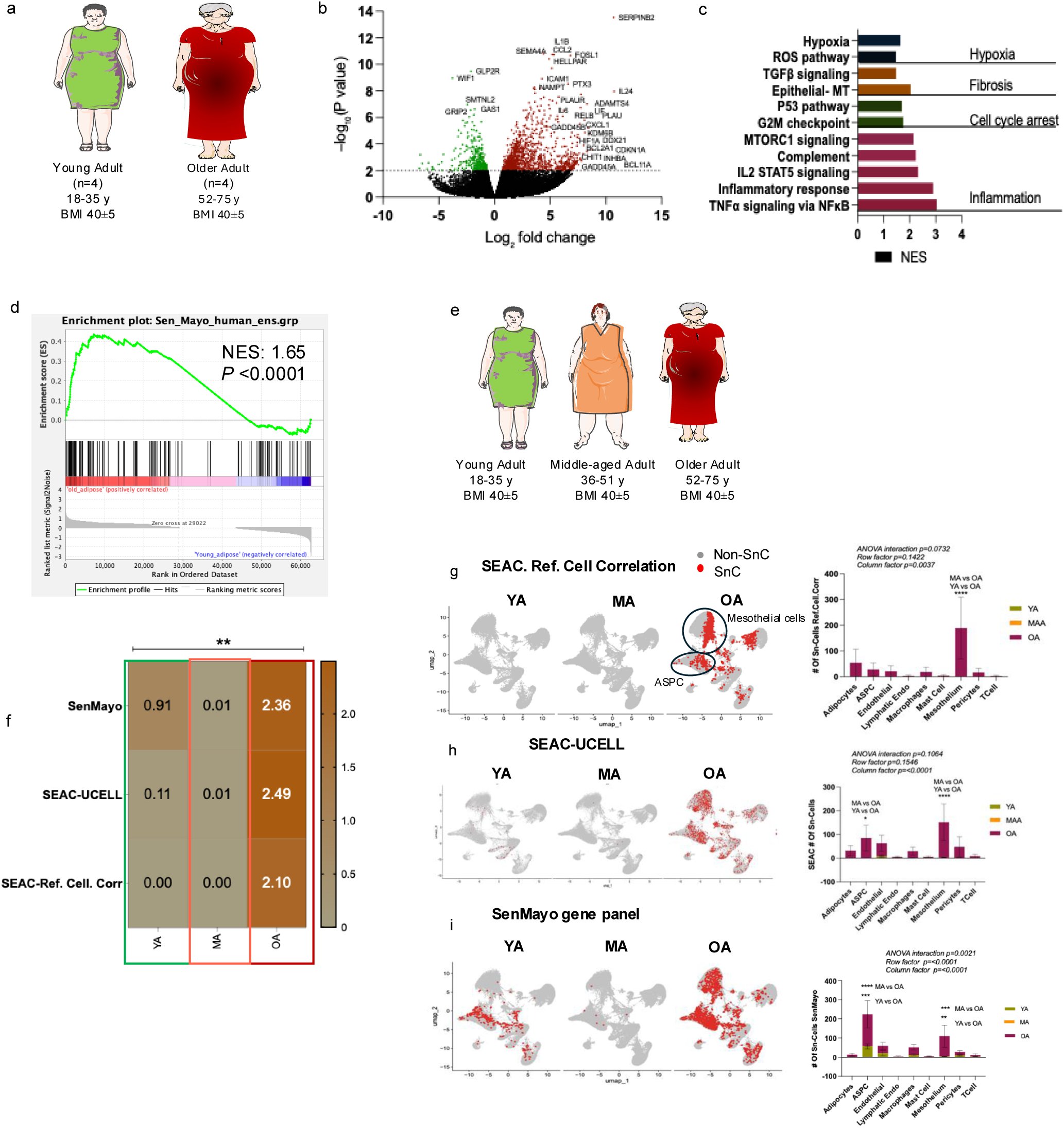
Transcriptomics analyses reveal increased cellular senescence in older obese adults than BMI-matched younger adults. **(a)** Sources of visceral adipose tissue (VAT) used to obtain Bulk-RNAseq data used in panels b-d. **(b)** Volcano plot indicating upregulation of senescence-associated biomarkers in older obese adult VAT relative to young. **(c)** GSEA indicating enrichment of pathways associated with canonical features of cellular senescence in older obese adult VAT compared to young (NES is normalized enrichment score). **(d)** NES plot indicating that VAT from older obese adults was positively correlated with the SenMayo gene panel, while younger obese adults VAT was negatively correlated. **(e)** Sources of VAT used to obtain snRNAseq data used in panels f-i. **(f)** Three different methods for predicting senescent cells (SnCs): SEAC-UCELL gene expression patterns, SEAC-reference cell correlation, and SenMayo gene set enrichment yielded a similar fraction of SnCs (0.01-2.49%) in human VAT per age-group. **(g-i)** All three SnC identification approaches revealed increased SnC burden in older obese adults (OA) compared to middle-aged (MA) and younger, predominantly in ASPCs and mesothelial cell populations. Individual donors were treated as independent biological replicates, rather than pooling nuclei across donors. Statistical significance was assessed using two-way ANOVA followed by Tukey’s post hoc multiple-comparisons test (age effect, *P = 0.0074*). Significance levels are indicated as P < 0.05 (*), P < 0.01 (**), P < 0.001 (***), and P < 0.0001 (****); ns, not significant.

We next performed single-nucleus RNA sequencing (snRNA-seq) to resolve age-associated changes at the cellular level. Because mature adipocytes are fragile and challenging to capture using conventional single-cell RNA sequencing, we used a single-nucleus approach. In addition to young and older adults, a middle-aged group to capture potential intermediate age-associated changes was included **(Fig. 1e).** UMAP analysis identified nine major cell populations **(Extended Fig. 2a)**, including adipocytes, mesothelial cells, adipose stem/progenitor cells (ASPCs), endothelial cells, lymphatic endothelial cells, pericytes, macrophages, T cells and mast cells. Cell-composition analysis revealed trends toward increased proportions of mesothelial and immune cells and decreased proportions of adipocytes, endothelial cells and pericytes in older obese adults compared with younger adults **(Extended Fig. 2b).** Cell-type and subcluster annotations were guided by the human adipose tissue atlas of Emont *et al.* (2022)^8^, using established lineage-specific marker genes **(Extended Fig. 2c-h).**

Sub-clustering further revealed age-associated expansion of specific cellular subsets, including VA3 adipocytes, hASPC6 progenitors and hMES1 mesothelial cells in older adults **(Extended Fig. 2i-l).** Notably, most endothelial subclusters were relatively enriched in middle-aged adults **(Extended Fig. 2l),** suggesting an age-dependent redistribution of the vascular compartment, with a potential transient enrichment during middle age that warrants further investigation.

We next identified SnCs within the VAT using three complementary approaches **(Supplementary Table 2).** First, we developed a SEAC-based approach comprising four senescence-associated features: **S**, SASP-associated genes; **E**, extracellular matrix (ECM)-related genes; **A**, anti-apoptotic genes; and **C**, cell-cycle-arrest genes. We selected genes within each SEAC category based on our bulk RNA-seq differential gene expression data and published literature^13,16,17^ and used these genes to calculate a SEAC-UCell score. Second, we generated a SEAC-defined senescent reference profile and identified cells exhibiting a Pearson correlation coefficient > 0.55 with this profile using the SEAC-Reference Cell Correlation approach. Third, we applied the SenMayo gene signature to further characterize senescence-associated transcriptional states and assess concordance across the three computational approaches.

All three approaches identified relatively small populations of predicted SnCs, comprising up to approximately 2.5% of the total VAT cell population, with SnC burden increasing with age **(Fig. 1f)**. Two-way ANOVA revealed a significant effect of age on senescence burden, whereas neither the method of SnC identification nor the method-by-age interaction was significant. These findings indicate a consistent age-associated increase in SnC burden across the three approaches, despite their distinct conceptual and computational frameworks.

Cell-type-resolved analysis further revealed selective vulnerability of specific cellular compartments to senesce in older obese adults. Two-way ANOVA with post-hoc analysis of the SEAC-based approach identified the ASPC and mesothelial compartments as particularly sensitive to age-associated increases in senescence **(Fig. 1g, h)**, while also demonstrating an overall increase in senescence across the tissue. Consistent with these findings, SenMayo analysis demonstrated significant increases in SnCs within ASPCs in young and old, but not middle-aged adults, as well as mesothelial cells only in the older adults **(Fig. 1i).** The Reference Cell Correlation identification method identified SnCs predominantly in the mesothelial population **(Fig. 1i)**, demonstrating good correlation amongst the various methods of identifying SnCs.

Senescence was not uniformly distributed across each cell-type but instead accumulated within discrete subpopulations, including VA3 and VA5 adipocytes, hASPC3 and hASPC6 progenitors, hMES1 mesothelial cells, hENDOS2 endothelial cells and hMAC3 macrophages and hMONO1 monocytes **(Extended Fig. 3a-o).** Importantly, in low BMI older adults, senescence accumulated in cellular subsets overlapping with those identified in the high-BMI cohort, including hASPC6, hMES1, hENDOS2 and VA3 **(Extended Fig. 4a-d).** This suggests that these particular cellular populations are susceptible to senescence independently of adiposity but as a characteristic of aging biology.

Finally, differential gene expression analysis of the snRNA-seq data using Seurat’s FindMarkers function showed 75% concordance with bulk RNA-seq for age-associated pathway perturbation **(Extended Fig. 5a, b)**, while MSigDB Hallmark pathway enrichment analysis identified 80% concordance of the pathways perturbed in VAT from older obese adults relative to young or middle-aged **(Extended Fig. 5c, d)** with no significant differences between MA and OA **(Extended Fig. 5e)**. The comparison of OA vs. YA and OA vs. MA produced a near identical rank order of the probability of pathways perturbed in older VAT relative to the two younger age groups **(Extended Fig. 5f)**. Collectively, these data demonstrate that aging is associated with increased SnC burden in VAT and that aging itself drives senescence more than obesity alone does. Senescence preferentially accumulated in ASPC and mesothelial cells, extending to adipocyte, endothelial and macrophage cell subsets. Furthermore, senescence-associated transcriptomic changes across age groups is non-linear, with middle age marking a transition into older age characterized by the consolidation and amplification of core SnC features.

### Enhanced intercellular communication centered on ASPCs and mesothelial cells in aged visceral adipose tissue

Next, we sought to explore how cell-cell communication changes with age in VAT in a setting of obesity. Intercellular communication levels were relatively comparable between young (YA) and middle-aged adults (MA) but increased markedly in older adults (OA) **(Fig. 2a)**, doubling in signal strength in old age. This was supported by MSigDB Hallmark pathway enrichment analysis of the top 200 differentially expressed genes from the YA *vs* OA, OA *vs* MA, and MA *vs* YA comparisons where MA and YA were equivalent and similarly different from OA **(Extended Fig. 5c-f)**.

**Figure 2.**
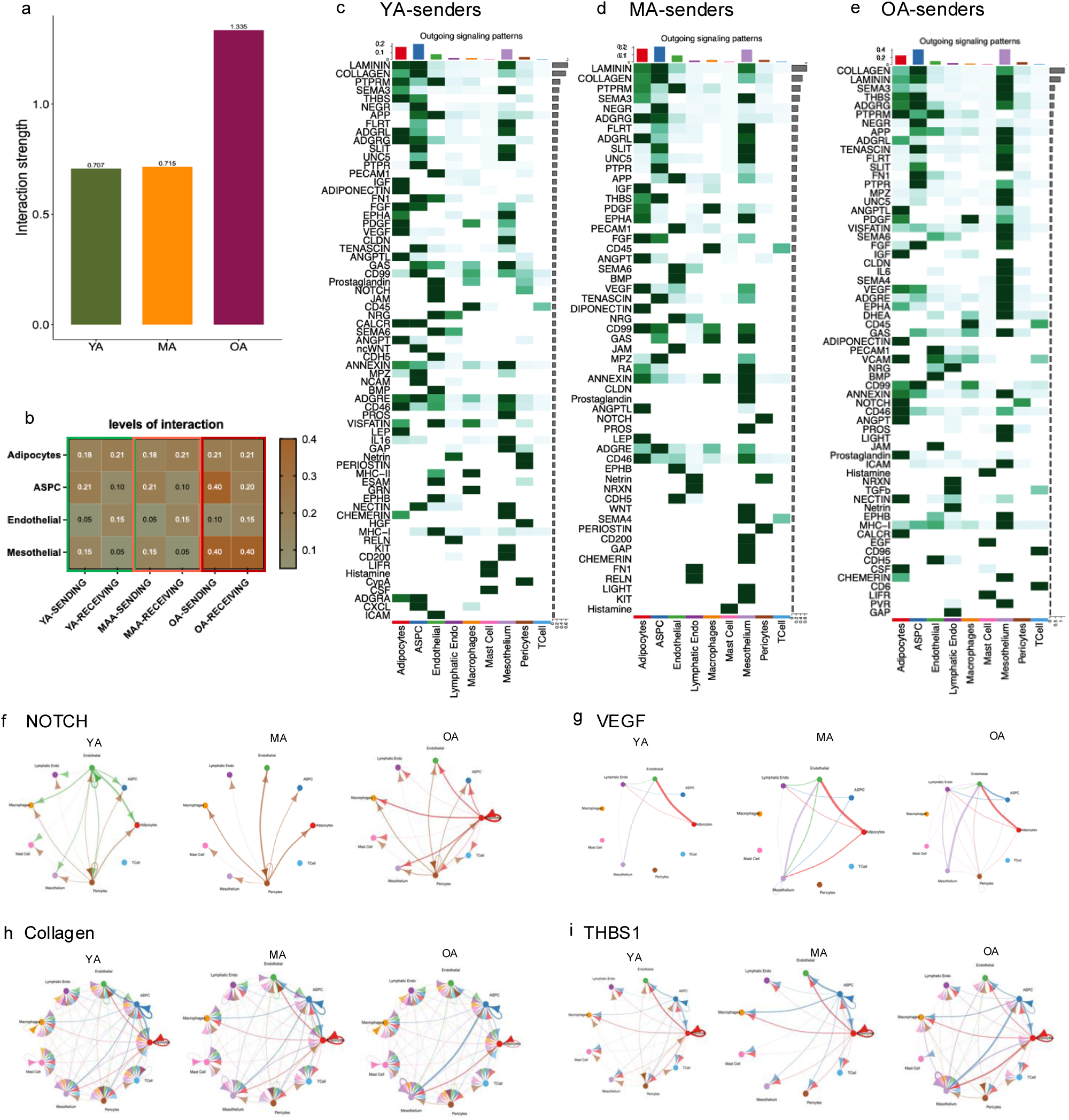
Senescent adipocyte stem/progenitor and mesothelial cells become dominant signaling hubs during aging. **(a)** CellChat analysis of snRNAseq data predicted an age-related increase in cell communications in the VAT from older obese individuals compared to MA or YA. **(b)** Senescent ASPCs and mesothelial cells from older adult VAT had the strongest outgoing signals, while mesothelial cells showed equivalent levels of outgoing and incoming signals. **(c-e)** Heatmaps indicating the strength of the outgoing signals by pathway (y-axis) and cell-type (x-axis); colors are used to indicate distinct cell-types. Histogram above the heatmap illustrates the total predicted amount of outgoing and incoming signal by cell-type. Shades of green are used to indicate the strength of the signal by cell-type. Grey bars to the right of the heatmap indicate the relative strength of that signal across all cell-types. Note the y-axes for OA is 2x greater than YA and MA. Older adults showed increased inflammatory and immune-modulatory cell-cell communication, represented by IL6, TGFβ, EGF, VCAM, and CD6 (red boxes), relative to younger individuals. (**f-i**) Circular plots showing the ligand-receptor signaling pathways predicted to be altered in VAT of older adults.

In older VAT, ASPCs and mesothelial cells, which were identified as senescence-enriched populations **(Fig. 1g-i)**, exhibited a high density of predicted outgoing signals also with a high density of incoming signals to mesothelial cells **(Fig. 2b)**. This suggests that ASPCs and mesothelial cells may serve as communication hubs within VAT, driven by age, not obesity. Heat-map analysis further revealed age-associated changes in key inflammatory and immunomodulatory pathways, including IL-6, TGF-β, EGF, VCAM, and CD6 **(Fig. 2c-e; Extended Fig. 6a-c)**. Further, pathways involved in tissue remodeling and inflammation, including NOTCH, VEGF, collagen, and THBS1 **(Fig. 2f-i)**, CSF, MHC-I, and ICAM, were altered in older adults **(Extended Fig. 6d-f)**, suggesting that aging is accompanied by a shift toward distinct intercellular signaling programs rather than a uniform increase in baseline communication.

### Senescent cells establish a conserved intercellular communication program

The patterns of outgoing and incoming signals from and to non-senescent and SnCs in VAT were analyzed using CellChat across the three methods used to identify SnCs.

### SEAC-Reference-Cell Correlation Approach

CellChat analysis based on SnCs identified by the SEAC-reference-cell correlation identified an outgoing signaling module (Pattern 4) shared by multi-lineage SnCs and non-senescent mast cells. This network included inflammatory and immune pathways (IL1, MIF, CSF3, and HISTAMINE), growth and morphogen-associated signals (EGF and WNT) and tissue-remodeling and stromal-associated mediators (PLAU, CYPA, and THY1) **(Fig. 3a)**. Predicted incoming signals were organized into three major patterns **(Fig. 3b)**. SnCs of multiple lineages collectively shared Pattern 5 and received signals that overlapped with components of their outgoing network, potentially indicative of autocrine and/or paracrine reinforcement of the senescent state. Non-senescent mesothelial cells formed Pattern 1 and preferentially received extracellular matrix, adhesion, inflammatory, and tissue-remodeling signals, including FN1, TENASCIN, VCAM, IL6, and PLAU. Mature adipocytes formed Pattern 2 and preferentially received endocrine, metabolic, and morphogen-associated signals, including TGFβ, WNT, ADIPONECTIN, DHEA, and TESTOSTERONE, suggestive of tissue remodeling and metabolic regulation.

**Figure 3.**
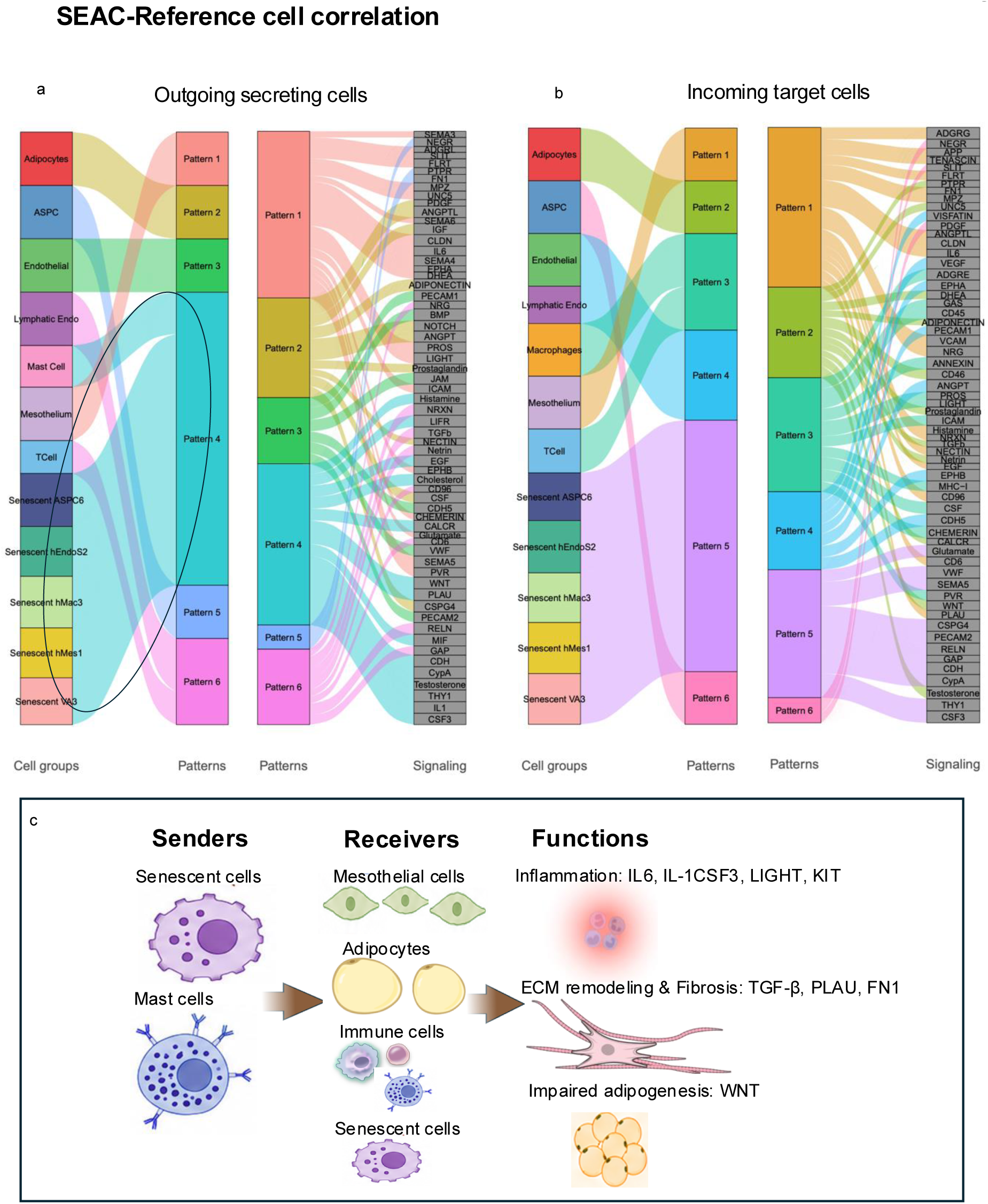
Senescent cells establish a conserved intercellular communication program. (a,c) CellChat Pattern Recognition Analysis of the snRNAseq data was used to identify recurring profiles of outgoing and incoming signals (patterns) by cell-type using the SEAC-Reference cell correlation to identify SnCs. In the left-most column are indicated signals from non-senescent cell-lineages; below are subclusters of cell-types enriched in SnCs (Endo/endothelia; Mac/macrophage; Mes/mesothelial; VA/visceral adipocyte). To the right (grey boxes) indicate the genes in each of the signaling patterns. **(a)** Outgoing and **(b)** incoming signaling interactions were each organized into six distinct communication patterns (distinct for outgoing and incoming). Similar communications analysis for the other two methods of SnC identification is provided in Extended Fig. 5. **(c)** Schematic representation of the common CellChat communication patterns identified using the SenMayo, SEAC-Reference Cell Correlation and SEAC-UCELL approaches to identify SnCs.

### SenMayo Gene Panel Approach

CellChat analysis of SenMayo-defined SnCs also predicted the same, shared outgoing communication network Pattern 5 emanating from senescent ASPCs, endothelial cells, macrophages, mesothelial cells and adipocytes **(Extended Fig. 7a)**. As described above, this pattern of predicted outgoing communications was shared with non-senescent mast cells, and included inflammatory and immune pathways (IL1, MIF, CSF3, and HISTAMINE), growth and tissue-remodeling pathways (EGF, PLAU, and WNT), and stromal-associated signals (CYPA and THY1). These data reinforce the evidence for shared secretions from SnCs across multiple lineages.

Unlike outgoing signaling, predicted incoming communication remained largely cell type-specific **(Extended Fig. 7b)**. Mesothelial cells, including senescent hMes1, preferentially received extracellular matrix remodeling, inflammatory, and adhesion-related signals, including TENASCIN, FN1, VCAM, IL6, and PLAU. Mature adipocytes preferentially received metabolic and morphogenetic signals, including ADIPONECTIN, WNT, TGFβ, and NETRIN, whereas macrophages, including senescent hMAC3, preferentially received myeloid-associated signals such as CSF, CSF3, and CHEMERIN.

Senescent adipocytes (VA3) showed a particularly distinct age-related shift in incoming communication compared with other mature adipocyte populations. VA3 exhibited reduced association with canonical metabolic signals, including ADIPONECTIN and DHEA, and increased association with immune-related pathways, including ICAM, MHC-I, PVR, and IL1 (Pattern 6). This shift suggests remodeling of senescent adipocyte communication toward enhanced immune-adipose crosstalk. Notably, SenMayo-positive hASPC6 and hENDOS2 did not exhibit detectable incoming signaling above the applied CellChat threshold. This finding should not be interpreted as an absence of incoming communication, as weaker interactions may have remained below the predefined threshold **(Extended Fig. 7b)**.

In contrast, non-senescent ASPCs maintained, across age-groups, bidirectional communication through progenitor– and extracellular matrix-associated pathways, including NEGR, FN1, PTPR, and CD34, while receiving trophic and migratory signals through PDGF, SLIT, and NEGR **(Extended Fig. 7b)**. Similarly, non-senescent endothelial cells maintained bidirectional communication through pathways associated with vascular remodeling (VEGFA, ANGPT, EPHA, and EPHB), vascular integrity and adhesion (PECAM and CDH5), and metabolic/inflammatory signaling (VISFATIN).

### SEAC-UCELL Approach

The SEAC-UCELL approach of identifying senescent cells revealed outgoing communication patterns broadly consistent with those identified by the SenMayo and SEAC-reference-cell correlation approaches. Most SnC types converged on a common outgoing signaling pattern (Pattern 6, **Extended Fig. 7c**), further supporting the emergence of a shared senescence-associated communication program across cell lineages. sn-hASPC6 was a notable exception, retaining an outgoing communication pattern more closely aligned with its parent ASPC lineage. SEAC-UCell-defined SnC populations did not emerge as distinct major signal recipients above the applied CellChat threshold **(Extended Fig. 7d)**. The exception was sn-hMES1, which retained an incoming signaling pattern similar to that of its parent mesothelial lineage. These findings suggest that senescence may preferentially reshape outgoing communication, whereas incoming signaling remains more strongly influenced by lineage identity. However, weaker incoming interactions below the applied CellChat threshold cannot be excluded.

Despite differences in how SnCs were identified, all three approaches converged on a common feature: SnCs across multiple lineages were predicted to have a coordinated outgoing communication program. This suggests that acquisition of a senescent state may impose a shared communication phenotype across otherwise transcriptionally and functionally distinct cell types. This is consistent with the robust SASP phenotype of SnCs but contradicts the notion that SnCs and their SASP is highly heterogenous based on cell lineage, senescence inducer, and tissue microenvironment, raising the possibility that the latter is an in vitro artifact.

In contrast, predicted incoming signals to SnCs were more heterogeneous and frequently retained lineage-specific features, which may be indicative of signals necessary to retain cell identity or viability. The predicted signaling networks further suggest that senescent cells impact adipose tissue function via driving chronic inflammation, altered immune-adipocyte communication, extracellular matrix remodeling and impaired adipogenesis. The SEAC-reference-cell correlation analysis additionally identified potential autocrine/paracrine signaling in senescent cells, suggesting a mechanism by which senescence could be reinforced or maintained.

### snRNAseq predicts senescence impairs adipogenesis and angiogenesis while augmenting inflammation and tissue remodeling in older adult VAT

To investigate the predicted functional consequences of cellular senescence in aging VAT, we integrated differential gene expression, RNA velocity, pathway enrichment, histological validation, and qPCR analysis across SnC populations. CellChat identified WNT9A-LRP6 signaling as a prominent interaction associated with SnCs **(Extended Fig. 8a-c)**. *WNT9A* was highly expressed in senescent populations across multiple adipose cell lineages, especially in VA3 compared with their non-senescent counterparts. In contrast, expression of *PPARG*, *ADIPOQ*, *PLIN1*, *FABP4*, *LIPE* and other key regulators of adipocyte differentiation and maintenance, was reduced in senescent cells **(Fig. 4a)**. These data are consistent with senescence impairing differentiation of ASPCs to mature adipocytes. This was supported by RNA velocity analysis **(Fig. 4b)**. ASPC6, a committed adipocyte progenitor population from which adipocytes are derived, exhibited a substantial senescence burden **(Extended Fig. 3a-c)**, resulting in reduced contributions towards visceral adipocytes.

**Figure 4.**
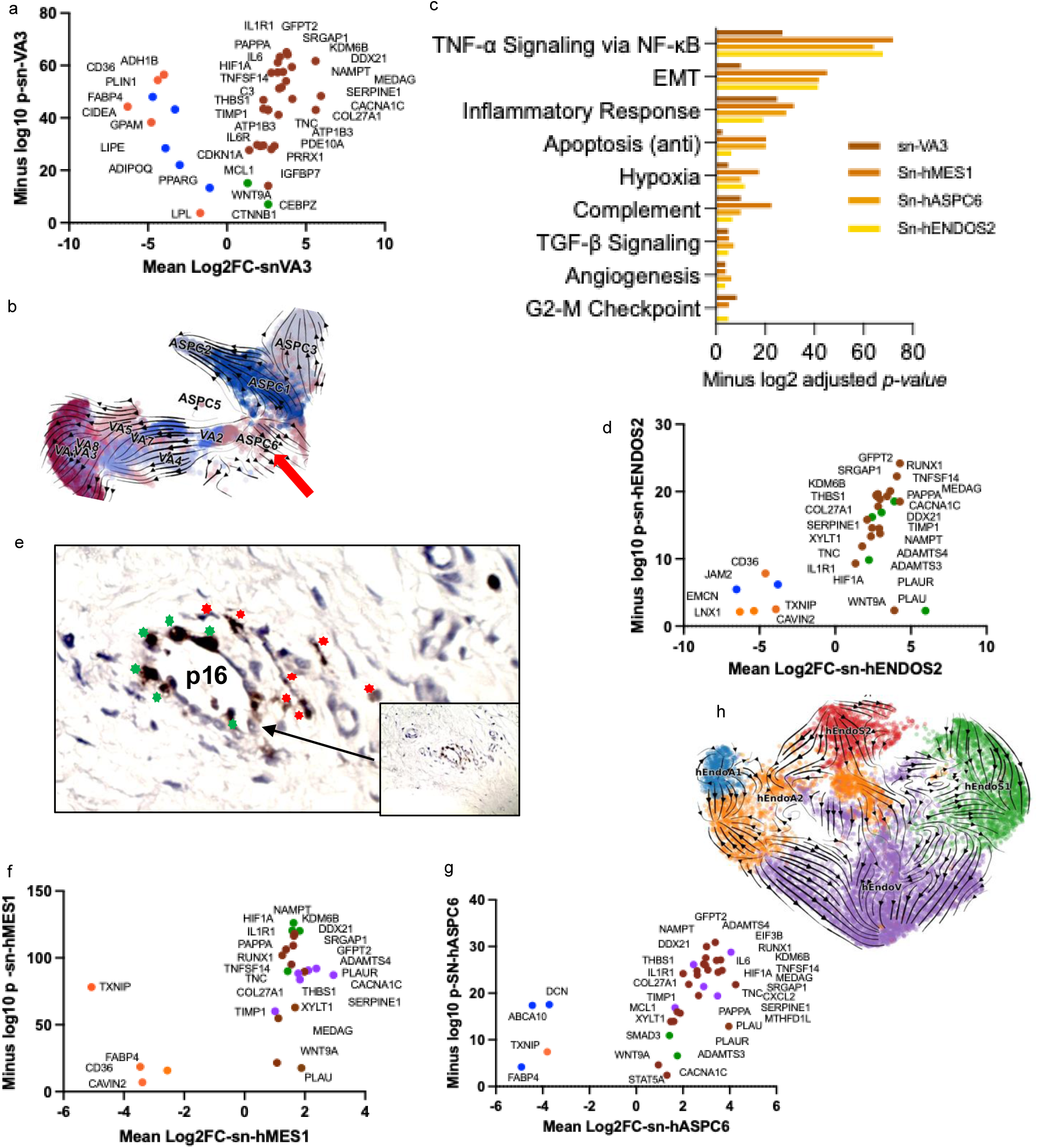
Senescent cell signaling in VAT impacts adipogenesis, inflammation and fibrosis. **(a)** Differential gene expression analysis (DGE) of snRNAseq data comparing senescent vs. non-senescent VAT-derived visceral adipocytes (VA) indicated increased expression of *WNT9A* and *CTNNB1* in senescent VA3, compared to its non-senescent VA (green dots). In contrast, key genes involved in adipocyte differentiation and maintenance, including *PPARG, ADIPOQ, PLIN1, FABP4,* and *LIPE*, were downregulated in this senotype (blue dots). **(b)** RNA velocity analysis on snRNA-seq of older obese adult VAT identified ASPC6, a committed adipocyte progenitor population, as a population with a substantial senescence burden. **(c)** Hallmark MSigDB pathway analysis of the top 200 differentially expressed genes across SnC populations in older obese adults revealed significant enrichment of TNF-α signaling via NF-κB, epithelial-mesenchymal transition, inflammatory response, apoptosis, hypoxia and complement, indicating a conserved inflammatory and tissue-remodeling program in four VAT cell types. **(d)** DGE of senescent human endothelial subcluster 2 *vs.* non-senescent endothelial cells exhibited reduced expression of cell-lineage markers, including *JAM2* and *EMCN* (blue dots, accompanied by increased expression of extracellular matrix remodeling– and fibrosis-associated genes, *THBS1*, *PLAU*, *PLAUR*, *COL27A*, *ADAMTS4*, and *TIMP1*, compared to non-senescent endothelial cells in VAT (green dots). **(e)** Immunohistochemical staining for p16^INK4a^ identified p16^+^ endothelial cells (green) within fibrotic vascular structures, as well as p16^+^ mesenchymal-appearing cells (red). **(f)** DEG between senescent human VAT mesothelial cells subcluster 1 and non-senescent hMES1 identified increased gene expression of inflammatory or SASP markers such as *TNFSF14*, *IL1R1*, *KDM6B*, and *NAMPT* (green dots), and ECM remodeling /fibrosis genes such as *ADAMTS4*, *PLAUR*, *SERPINE1*, *TIMP1*, *TNC* and *COL27A* (purple dots). **(g)** DGE analysis of senescent hASPC subcluster 6 cells revealed significant upregulation of *WNT9A* and *SMAD3* (green dots) and downregulation of *FABP4*, *DCN*, and *ABCA10* (blue dots) relative to non-senescent ASPCs, suggesting altered adipogenic regulation. Concurrent upregulation of ECM remodeling– and fibrosis-associated genes, including *THBS1*, *COL27A1*, *TIMP1*, *SERPINE1*, and *ADAMTS4* (purple dots), further suggests a shift toward a profibrotic transcriptional program. **(h)** RNA velocity analysis of endothelial subpopulations identified hENDOS2 (red) as a senescence-enriched population with predicted transitions toward other endothelial states, suggesting that senescence in this progenitor-like endothelial population may contribute to impaired vascular remodeling and angiogenesis.

To define broader transcriptional consequences of senescence, we performed Hallmark MSigDB analysis using the top 200 differentially expressed genes from senescent versus non-senescent populations for each cell-type. Senescent adipocytes, mesothelial cells, ASPCs, and endothelial cells showed prominent enrichment of TNF-α signaling via NF-κB, inflammatory response, epithelial-mesenchymal transition, complement, hypoxia, and apoptosis-related pathways **(Fig. 4c)**. This reinforces the notion that SnCs across adipose cell lineages acquire a shared inflammatory and cellular-stress phenotype accompanied by tissue-remodeling signatures.

Senescent hENDOS2 cells exhibited reduced expression of the endothelial identity markers JAM2, EMCN, etc., with increased expression of ECM remodeling/fibrosis-associated genes, including THBS1, PLAU, PLAUR, COL27A, ADAMTS4, TIMP1 **(Fig. 4d)**, suggestive of a loss of endothelial identity and acquisition of a remodeling-associated phenotype. Immunohistochemistry provided evidence of senescence within remodeled vascular structures. p16^+^ endothelial cells were detected within fibrotic small venous structures, together with p16^+^ mesenchymal-appearing cells **(Fig. 4e)**, linking senescence endothelial cells to a fibrotic vascular microenvironment.

Across senescent cell-types, many ECM remodeling/fibrosis-related genes, inflammatory, and senescence-associated gene expression were increased relative to non-SnCs **(Fig. 4 a,d,f,g).** In addition, WNT-related genes, as well as *KDM6B*/JMJD3, an epigenetic histone demethylase implicated in adipogenic, inflammatory, and senescence-associated programs, were more highly expressed in VAT SnCs **(Fig. 4 a,d,f,g)**.

RNA velocity analysis identified hENDOS2 as a likely source of several other endothelial cell subpopulations **(Fig. 4h)** Senescence of hENDOS2 cells, with loss of cell identity and an increased remodeling-associated phenotype may compromise normal endothelial state transitions and contribute to impaired vascular remodeling and angiogenic capacity. Notably, again, these phenotypes are driven by age-related senescence instead of obesity.

Finally, to determine whether the inflammatory signature identified computationally was evident at the tissue level, we measured TNF expression by qPCR in VAT across age groups. TNF expression was increased in VAT from older adults compared with younger and middle-aged groups **(Extended Fig. 8d),** providing independent tissue-level support for increased TNF-associated inflammatory signaling with aging. We next measured the expression of p21/CDKN1A **(Extended Fig. 8e),** and p16/CDKN2A **(Extended Fig. 8f)** and canonical markers of cellular senescence, by qPCR. Both markers were increased in VAT from older adults, further supporting the increased senescence burden associated with aging.

### Conserved transcriptional programs support CellChat-predicted functional consequences of multicellular senescence and identify KDM6B as a candidate epigenetic regulator

To further characterize the regulatory programs associated with cellular senescence, we performed decoupleR transcription factor activity analysis comparing senescent and non-SnCs across lineages. SnCs showed increased inferred mean activity of multiple stress-, inflammatory-, hypoxia-, and tissue-remodeling-associated regulators, including NF-κB, AP-1, JUN, FOS, RELA, STAT3, EPAS1, HIF1A, TP53, SP3, PRRX1, SP7, NFATC1, SMAD2, SMAD3, RUNX2, TCF7L2, and CTNNB1 **(Fig. 5a, b)**. Notably, PRRX1 showed the highest inferred mean activity in senescent hENDOS2 cells, consistent with activation of a mesenchymal/tissue-remodeling program.

**Figure 5.**
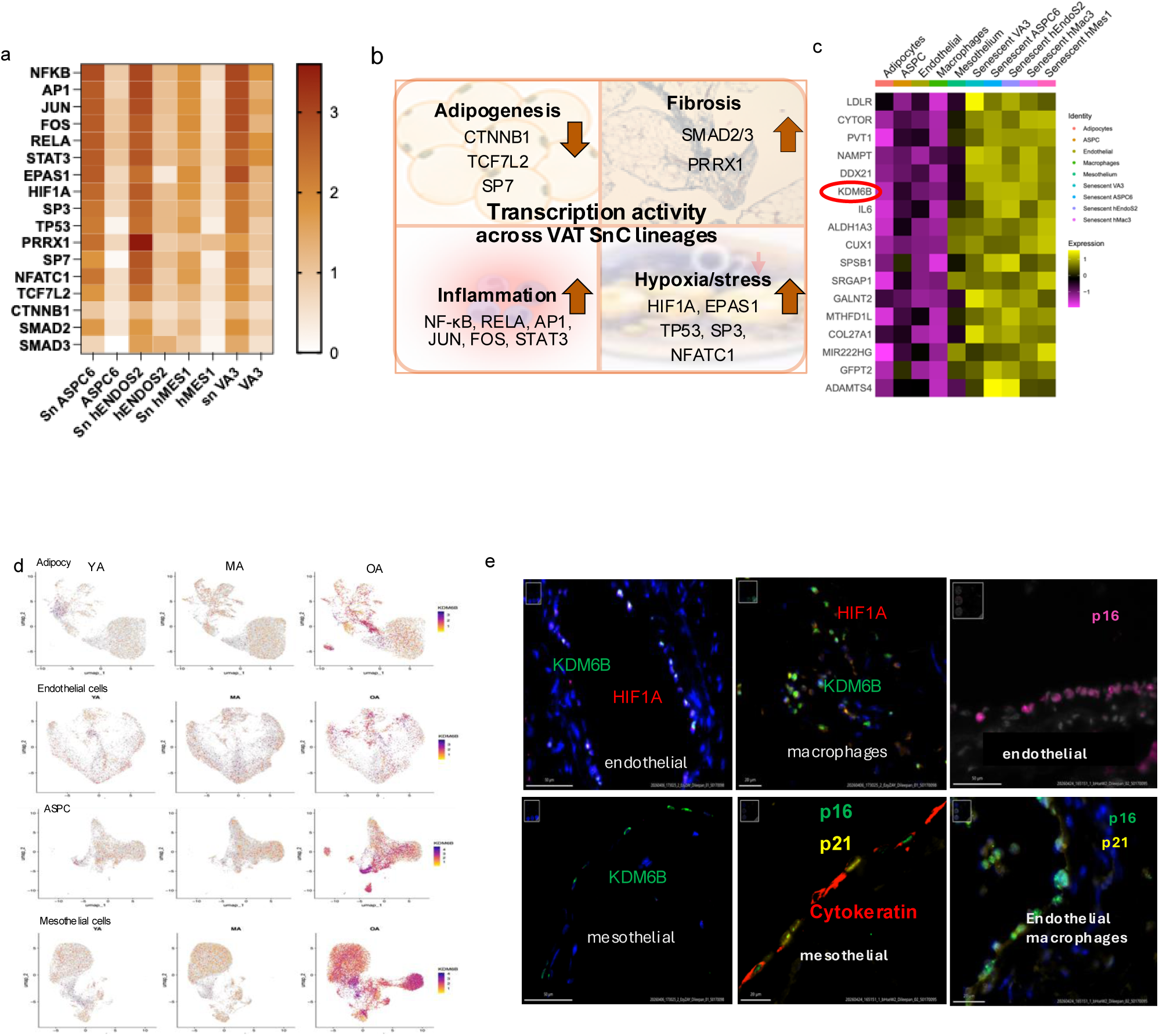
Conserved transcriptional programs support the CellChat-predicted functional consequences of multiple senescent cell types in VAT and identify *KDM6B* as a candidate epigenetic regulator. (a) DecoupleR-inferred mean transcription factor activity in senescent *vs.* non-senescent cells across lineages from the snRNAseq data, identified stress, inflammatory, hypoxia and tissue-remodeling-associated regulators. (b) Schematic summarizing the functional outcomes identified by transcriptional programs and CellChat analysis. (c) Heatmap showing the shared relative expression of genes including *KDM6B*, in SnCs across all lineages relative to their non-senescent counterparts. (d) UMAPs showing *KDMcB* expression across cell types and age groups. (e) Representative COMET proteomics images showing KDM6B and HIF1A expression in endothelial cells, macrophages, and mesothelial cells, as well as p16^INK4A^
and p21^CIP1^
expression in endothelial cells (CD31^+^), macrophages (CD68^+^) and mesothelial (pan-cytokeratin^+^) cells.

The enrichment of HIF1A and EPAS1 as well as TP53-associated stress responses suggest that age-related senescence in VAT may be induced by hypoxia and genotoxic stress **(Fig. 5b).** Among the shared differentially expressed genes identified across predicted SnC lineages **(Fig. 5c)**, we prioritized KDM6B, an H3K27me3 demethylase, as a candidate epigenetic regulator because of its established roles in cellular senescence, inflammatory signaling, and adipose cell differentiation^18–22^. Its differential expression across multiple SnC populations suggests a potential role in coordinating senescence-associated transcriptional programs and tissue remodeling in aged VAT. UMAP analysis confirmed strong age-dependent up-regulation of KDM6B expression across cell-types demonstrated to senescence in VAT **(Fig. 5d)**, which rises sharply after middle-age. This was confirmed at the protein level in VAT from older obese adults **(Fig. 5e, Extended Fig. 8k-m)** together with enhanced TP53-associated activity suggesting a potential link between stress-induced epigenetic remodeling and the establishment or maintenance of the senescent state. Visium HD spatial transcriptomics also supported increased expression of *KDM6B* across cell types in VAT from old obese individuals relative to young **(Extended Fig. 8n-s)** as well as increased expression of *p21/CDKN1A*, while *p16/CDKN2A* expression was increased only in adipocytes and whole VAT, but not in the other cell-types. KDM6B expression was predominantly localized to mesothelial and endothelial cells **(Extended Fig. 8p, q)**.

## Discussion

Aging and obesity are both associated with cellular senescence, progressive metabolic dysfunction, and implicated as drivers of many age-related chronic diseases. A high fat diet is often used to induce obesity in preclinical models to drive accelerated cellular senescence. However, the question of whether obesity or aging drives senescence more robustly is not well explored in humans. Here, we demonstrate the sustained obesity through middle-age (≤50 years old) is not associated with senescence in the VAT to nearly the extent as what occurs in old age. This suggests that chronological age is a greater risk factor for senescence than obesity. This may also imply that obesity drives age-related diseases through mechanical and metabolic stress rather than through cellular senescence, which is established for aging.

Epidemiological studies demonstrate that excess adiposity is associated with increased lifetime cardiovascular risk, heart failure and mortality, while recent longitudinal evidence indicates that long-term obesity is associated with features of accelerated biological aging even in relatively young adults^23–25^. Correa-Burrows *et al.* showed that persistent obesity in young adults is associated with epigenetic age acceleration and other markers of biological aging, supporting the concept that obesity accelerates age-related processes. Furthermore, accelerated biological aging is associated with epigenetic age acceleration, telomere shortening, and chronic inflammation, suggesting a potential contribution of cellular senescence. By examining BMI-matched human visceral adipose tissue from young, middle-aged and older adults, and integrating single-nucleus transcriptomics, multiple senescence-detection strategies, cell-cell communication, transcription-factor activity and spatial proteomics, we identify a convergent multicell-lineage senescence program capable of driving organ dysfunction that only becomes prominent only in older adults.

Adipose tissue is increasingly recognized as a highly dynamic and heterogeneous organ in which aging affects cell types and depots differentially ^8,26, 27^. The greatest remodeling of adipose composition occurs in middle-age, particularly in the ASPC compartment ^27^. We extend these results under conditions of sustained obesity. Older adults (in this case >50-years-old) exhibited substantially greater SnC burden than those ≤50-years-old. Pseudo-bulk pathway analysis revealed extensive overlap in pathways altered between older and young or middle-aged adults, but few differences between young and middle-aged obese adults. Thus, rather than senescence in VAT occurring across ages of obese individuals, or a progressive increase in senescent cell burden with age, senescence in obesity is driven by older age.

A strength of our study is the use of three strategies to identify SnCs within complex human tissues. SenMayo is a widely used SnC gene signature that emphasizes inflammatory and stress associated markers^13^. However, it does not include the canonical cell-cycle arrest markers p16^INK4a^ (CDKN2A) and p21^CIP1^ (CDKN1A), and inflammatory and stress-associated programs can also occur independently of cellular senescence. Thus, SenMayo-based scores may be influenced by the inflammatory state of cells, particularly in tissues such as adipose tissue. More broadly, the heterogeneity and tissue dependence of senescence-associated phenotypes highlight the limitations of relying on a single transcriptional signature to define senescence. To provide a complementary, multidimensional assessment of senescence, we designed the SEAC framework to identify SnCs based on four functional features of senescence: SASP, extracellular matrix (ECM) remodeling, anti-apoptotic signaling, and cell-cycle arrest. We complemented this with reference-cell correlation and a curated bulk/literature-derived senescence signature. These three approaches provided a highly convergent prediction of the senescent landscape in human VAT. All three methods predicted age-related senescence predominantly in ASPCs and mesothelial cells. SEAC and reference cell, not trained on inflammation alone, indicated a broader distribution of senescence more multiple cell lineages while continuing to highlight ASPCs and mesothelial cells as prominent populations.

The identification of discrete senescence-enriched populations is consistent with recent single-nucleus and spatial analyses showing selective vulnerability of specific adipose cell-types to stress and senescence. Miranda *et al.* identified increased susceptibility to senescence among adipocytes, macrophages, ASPCs, pericytes and endothelial cells in human subcutaneous adipose tissue ^28^. In VAT, we find ASPCs, mesothelial and endothelial cells have the greatest senescence in older obese adults, with the greatest predicted functional consequences on tissue function. In both depots, mature healthy adipocytes are depleted in older adults.

The identification of senescent mesothelial cells is notable because mesothelial cells are increasingly recognized as multifunctional stromal populations involved in tissue organization, extracellular-matrix remodeling and intercellular signaling. Their high senescence burden in our study, together with their prominent outgoing/incoming signaling activity, suggests that mesothelial senescence may have effects beyond the mesothelial compartment itself and impacting the microenvironment of multiple cell types In VAT with aging. Importantly, senescent mesothelial cells may also metastasis in older adults.

CellChat analyses across the three senescence-identification approaches revealed both shared and lineage-specific cell communication patterns. Outgoing signaling largely converged on a common transcriptional pattern across senescent cells of multiple lineages. This suggests that *in vivo* there may be common senotypes even across multiple cell-types. This predicted secretory pattern of senescent cells is also observed in non-senescent mast cells, suggesting outgoing signals from SnCs activate the innate immune system.

In contrast, incoming signals to SnCs are more heterogeneous. SenMayo revealed lineage-specific incoming signals. SEAC-UCELL identified limited incoming signaling to senescent cells. SEAC-Reference-cell correlation suggested preferential signaling within the senescent cell compartment, potentially reflecting autocrine/paracrine reinforcement of the senescent state. The method-dependent variation in incoming communication may reflect both analytical differences in SnC identification and biological heterogeneity among senescence-associated states, rather than differences in sensitivity alone. SnC signaling is predicted to target non-senescent mesothelial cells impinging on ECM-remodeling and pro-fibrotic pathways, including PLAU-PLAUR signaling, and to target adipocytes through WNT signaling predicted to negatively impact adipogenesis. Additional inflammatory signaling pathways, including IL1, IL6, and CSF3, suggest potential interactions between senescent and immune cells.

The predicted signaling changes unique to older adults were supported by transcription-factor activity and differential gene expression analyses. SnCs exhibited increased inferred activity of regulators associated with inflammatory signaling, cellular stress, hypoxia, extracellular-matrix remodeling and cell-fate regulation, including NF-κB, AP-1/JUN/FOS, STAT3, HIF1A, EPAS1, TP53, SMAD2/3, PRRX1, SP7 and RUNX2. The strong PRRX1 activity observed in senescent hENDOS2 cells suggests activation of a mesenchymal and tissue-remodeling program in this endothelial state, whereas the prominent SP7, TP53 and SMAD3 activity in hASPC6 is consistent with altered progenitor fate and adipogenic regulation.

These findings are consistent with previous studies showing that inflammatory and WNT/TGF-β-associated signals can impair adipocyte differentiation and alter stromal cell function in obesity. In human preadipocytes, inflammatory signaling involving WNT and TNF-α are linked to impaired differentiation^29^, while stromal populations can suppress adipogenesis through paracrine mechanisms^30^. PRRX1 was shown to inhibit adipogenesis through activation of TGF-β signaling^31^. Our findings also complement a recent human adipose atlas demonstrating extensive heterogeneity and metabolic states amongst mature adipocyte^7,32^. The distinct signaling phenotype identified in senescent VA3 cells may represent a state transition within an already heterogeneous adipocyte population.

Among the regulatory changes identified in our study, KDM6B emerged as a candidate epigenetic regulator potentially linking cellular senescence, inflammatory signaling, extracellular matrix remodeling, and impaired adipogenic function in visceral adipose tissue (VAT) from older adults with obesity. KDM6B, also known as JMJD3, is an inducible histone lysine demethylase that primarily removes repressive H3K27me3 marks, thereby influencing chromatin accessibility and transcriptional responses to inflammatory and cellular stress signals. Previous studies have implicated KDM6B in cellular senescence through regulation of the CDKN2A (p16INK4a) locus and interactions with p53-associated pathways^33^. In our study, KDM6B expression was associated with putative SnC populations exhibiting increased inferred activity of TP53, HIF1A/EPAS1, NF-κB/AP-1, and SMAD-associated transcriptional programs. These regulatory networks are involved in cellular stress responses, inflammatory signaling, extracellular matrix remodeling, and tissue fibrosis, suggesting that KDM6B may contribute to the coordinated transcriptional changes observed across senescence-associated cell populations. Consistent with this interpretation, increased KDM6B expression was accompanied by elevated expression of the senescence-associated markers CDKN2A (p16INK4a) and CDKN1A (p21CIP1) in VAT from older adults with obesity, although these associations do not establish a direct regulatory relationship.

Beyond its potential involvement in senescence-associated inflammatory programs, KDM6B has also been implicated in adipocyte differentiation through its regulation of chromatin – dependent transcriptional networks, including TGF-β-associated signaling. Previous studies have reported that suppression of KDM6B can promote adipogenic differentiation in specific experimental contexts^34^, suggesting a possible connection between KDM6B activity and the regulation of adipose progenitor cell fate. This relationship is particularly relevant to our findings of senescence-associated transcriptional remodeling in adipose stromal/progenitor cells (ASPCs), accompanied by altered signaling pathways potentially associated with impaired adipogenesis. Together, these observations raise the possibility that KDM6B contributes to the establishment or maintenance of a senescence-associated fibro-inflammatory microenvironment while influencing the adipogenic potential of resident progenitor cells. However, the functional consequences of KDM6B activity are likely to depend on cellular context, and its precise role in human visceral adipose tissue remains to be established.

Taken together, our findings support the conclusion that chronological aging drives cellular and molecular remodeling within the VAT that obesity alone does not produce. Despite the rarity of these cells, their rich intercellular signaling is predicted to have dramatic effects on the VAT in old age including driving impaired adipogenesis and angiogenesis, ECM remodeling, and activation of innate immunity. ASPCs and mesothelial cells emerge as particularly senescence-enriched populations and prominent signaling hubs, while senescent populations across lineages converge on a shared inflammatory and tissue-remodeling secretory program that is likely reinforced through autocrine and paracrine mechanisms.

### Novelty Statement

Our study reveals that age-associated senescence in human omental visceral adipose tissue, within the context of consistently high adiposity, is not simply characterized by the accumulation of putative SnCs but also by the emergence of a coordinated multicellular communication program. Despite their distinct cellular origins, predicted senescent cell populations converge on shared outgoing signaling patterns that may contribute to inflammation, extracellular matrix remodeling, and dysregulated adipogenesis within the aging visceral adipose

## Acknowledgement

We gratefully acknowledge the staff of the Clinical and Translational Science Institute (CTSI), particularly Adam Lewis, Jennie Brocksome, and Laurie Rachel, for their services, supported by NIH Award UM1TR004405.

## Funding

This study was supported by the National Institutes of Health (NIH) Common Fund (U54AG076041).

---

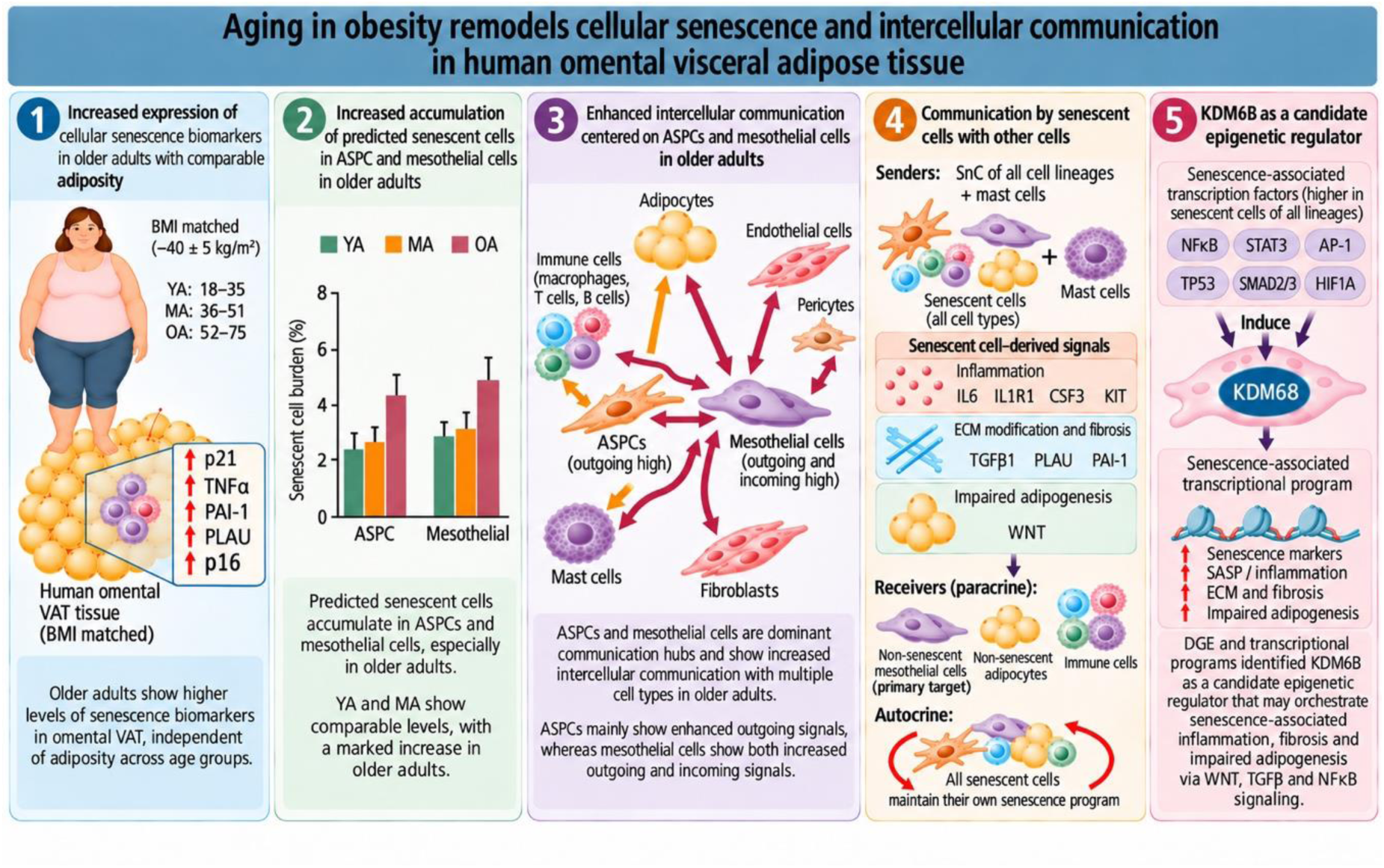

## Notes

### Competing Interest Statement

The authors have declared no competing interest.

