## Supplementary Tables for "Aging drives senescence in visceral adipose to a greater extent than obesity, instigating multicellular common signaling patterns that impair organ function and drive inflammation"

**Supplementary Table 1. Donor metadata**

| ID | YA1 | YA2 | YA3 | MA1 | MA2 | MA3 | MA4 | MA5 | OA1 | OA2 | OA3 | OA4 |
|---|---|---|---|---|---|---|---|---|---|---|---|---|
| AGE (years) | 25 | 26 | 27 | 41 | 42 | 43 | 50 | 51 | 64 | 65 | 68 | 71 |
| BMI (kg/m$^2$) | 36.3 | 40.1 | 39.48 | 38.73 | 37.89 | 42.51 | 40.41 | 39.58 | 43.84 | 41.1 | 35.4 | 37.2 |
| HOMA-IR | 7.1 | 3.64 | 3.5 | | 8.3 | 1.72 | 2.19 | 1.42 | 6.3 | NA | NA | NA |
| HbA1c (%) | 5.3 | 4.8 | 5.6 | 5 | 5.8 | 5.5 | 5.4 | 5.3 | 6.1 | 9.2 | NA | NA |
| Glucose (mg/dl) | 85 | 90 | 100 | 95 | 110 | 86 | 86 | 81 | 109 | NA | NA | NA |
| Insulin (mIU/L) | 33.8 | 16.4 | 14.2 | NA | 30.4 | 8.1 | 10.3 | 7.1 | 23.4 | NA | NA | NA |
| FFA_(mmol/) | 1.1 | 0.71 | 0.64 | NA | 0.6 | 0.73 | 1.05 | 0.9 | 0.89 | NA | NA | NA |
| Cholesterol (mg/dl) | 136 | 186 | 165 | 249 | 163 | 181 | 211 | 111 | 207 | 178 | NA | NA |
| HDL (mg/dl) | 23 | 55 | 43 | 37 | 40 | 47 | 55 | 39 | 55 | 38 | NA | NA |
| LDL mg/dl | 84 | 114 | 97 | 162 | 85 | 116 | 145 | 38 | 132 | 100 | NA | NA |
| TG (mg/dl) | 143 | 84 | 125 | 251 | 188 | 90 | 55 | 169 | 98 | NA | NA | NA |

**Supplementary Table 2. Approaches to identify senescent cells**

| SASP | EXTRA CELLULAR MATRIX | ANTI-APOPTOTIC | CELL CYCLE ARREST |
|---|---|---|---|
| IL1B+ | PLAUR+ | SERPINE1+ | CDKN1A+ |
| IL6+ | PLAU+ | SERPINEB2+ | TP53+ |
| CCL7+ | TNC+ | BCL2L1+ | GADD45A+ |
| CXCL8+ | VCAM1+ | GZMB- | CDKN2A+ |
| CCL2+ | ADAMTS4+ | RELT- | CCNB1- |
| CXCL2+ | TMEM217+ | MCL1+ | CCNB2- |
| CD55+ | ICAM4+ | | CDK1- |

**SEAC approach: S**- SASP, **E**- Extracellular matrix, **A**- Anti-apoptotic, **C**- Cell cycle arrest
Cells that simultaneously express SEAC-genes are considered senescent.
data.1 <- subset (data, Anti_Apoptopic_UCell > 0.2 & Cellcycle_related_UCell > 0.1 & Extracellular_Matrix_UCell > 0.1 & SASP_UCell > 0.1)

**SEAC-Reference-cell correlation**

Finding a reference cell in G1/G0 phase with expressing SEAC genes set at following threshold: CCL2 >1 **&** IL6 >1 **&** ICAM1 >1 **&** SERPINE1 >1 **&** MCL1 >1.5 **&** CDKN1A >1.5 & GADD45B >1 **&** MKI67 =0 **&** PCNA =0 **&** Phase = G1/G0
Cells that show greater than 55% similarity of the reference cell transcriptome profile is considered as senescent

**The SenMayo gene panel**

A UCell score threshold was determined using the R quantile () function (probs = 0.99) and applied to identify nuclei exhibiting enrichment of the 125-gene SenMayo senescence signature.

**Supplementary Table 3**. Detailed summary of CellChat-predicted outgoing and incoming signaling patterns, participating cell types, and associated ligand-receptor interactions.

| | **Outgoing signals** | **Outgoing Pattern** | **Incoming Pattern** | **Receiving signals** | **Direct ligand-receptor communication** | **Indirect communication** | **Functions** |
|---|---|---|---|---|---|---|---|
| **SenMayo** | SnC of all lineages | 5 | 1 | Mesothelium and sn-hMes1 | Histamine, EGF, PLAU, CypA & CDH | FN1, IL6, VCAM | Inflammation-IL6, VCAM, Histamine, CypA |
| | with mast cells | | | | | | ECM remodeling & fibrosis-PLAU, FN1 |
| | | | | | | | Stromal cell interactions-CDH |
| | | | 2 | Adipocytes | WNT, CADM, Testosterone | TGF-b | Impaired adipogenesis-WNT |
| | | | | | | | Fibrosis-TGF-b |
| | | | | | | | Cell adhesion-CADM |
| | | | | | | | Metabolism-testosterone |
| | | | 3 | Macrophages & sn-hMAC3 | CSF3 | Light, CSF, Chemerin | Inflammation-CSF3, LIGHT, chemerin |
| | | | 6 | Mast cell, Tcell and sn-VA3 | IL-1 | Glutamate, ICAM, KIT | Inflammation-glutamate, ICAM, KIT |

| | Outgoing signals | Outgoing Pattern | Incoming Pattern | Receiving signals | Direct ligand-receptor communication | Indirect communication | Functions |
|---|---|---|---|---|---|---|---|
| **SEAC-Ref.Cell. Correlation** | SnC of all lineages with mast cells | 4 | 1 | Mesothelium | Histamine, EGF, PLAU | FN1, IL6, VCAM | Inflammation-VCAM, Histamine<br>ECM remodeling & fibrosis-PLAU, FN1 |
| | | | 2 | Adipocytes | WNT, CALCR Testosterone | TGF-b | Impaired adipogenesis-WNT<br>Fibrosis-TGF-b<br>Cell adhesion-CADM<br>Metabolism-testosterone |
| | | | 5 | SnC of all lineages | CSF3, CDH, CypA, Thy1 | Glutamate, SEMA5 | Stromal cell interactions-CDH<br>Fibroblast-stromal signaling-THY1<br>Inflammation-glutamate, CSF3, CypA |

| | Outgoing signals | Outgoing Pattern | Incoming Pattern | Receiving signals | Direct ligand-receptor communication | Indirect communication | Functions |
|---|---|---|---|---|---|---|---|
| **SEAC-UCELL** | SnC of all lineages | 6 | 1 | Mesothelium & sn-hMes1 | Histamine, EGF, PLAU | FN1, IL6, VCAM | Inflammation-VCAM, Histamine |
| | with mast cells | | | | | | ECM remodeling & fibrosis-PLAU, FN1 |
| | | | 2 | Adipocytes | WNT, CADM, Testosterone | TGF-b | Impaired adipogenesis-WNT |
| | | | | | | | Fibrosis-TGF-b |
| | | | | | | | Cell adhesion-CADM |
| | | | | | | | Metabolism-testosterone |
| | | | 6 | Mast cells, T-cells | KIT, IL-1 | Glutamate, ICAM | Inflammation-IL-1, glutamate, ICAM, KIT |
