## Extended Figure Legends for "Aging drives senescence in visceral adipose to a greater extent than obesity, instigating multicellular common signaling patterns that impair organ function and drive inflammation"

**Extended Figure 1. Age-associated changes in signaling pathways. (a-l)** Gene set enrichment analysis (GSEA) of pathways identified as significantly upregulated in the visceral adipose tissue (VAT) of older obese adults compared to younger individuals with similar BMIs using BulkRNAseq data (n=4 per age group). Pathways identified included those associated with inflammation (TNF-α signaling via NF-κB, and the inflammatory response IL-6-JAK-STAT3 signaling, as well as complement), hypoxia and oxidative stress (hypoxia and reactive oxygen species), DNA damage and cell-cycle arrest (IFN-γ response, UV response up, G2M check point and p53 pathway) and fibrosis (TGF-β signaling and epithelial-mesenchymal transition). The top graph in each panel shows the ranked gene list based on the ranking metric, with genes ordered from the most positively to the most negatively associated. The bottom graph displays the running enrichment score (green curve) across the ranked gene list, while the black vertical ticks indicate the positions of genes belonging to the corresponding gene set. NES stands for Normalized Enrichment Score. It indicates the direction and relative strength of enrichment after normalization for gene-set size; q-value is False Discovery Rate (FDR)-adjusted significance measure. It accounts for multiple hypothesis testing when evaluating many gene sets or pathways.

**Extended Figure 2. Analysis of the snRNAseq data from obese adult VAT. (a)** UMAP analysis identified nine major cell populations in human VAT including adipocytes, mesothelial cells, adipose stem and progenitor cells (ASPCs), endothelial cells, pericytes, macrophages, T cells, mast cells, and lymphatic endothelial cells. **(b)** Cell-composition analysis revealed trends toward increased proportions of mesothelial and immune cells and decreased proportions of adipocytes, endothelial cells, and pericytes in older obese adults compared with younger adult VAT. **(c)** Expression of marker genes used to identify major cell-types. **(d-h)** Expression of marker genes used to identify subclusters of cell-types. **(i-l)** Sub-clustering of cell-types revealed age-associated expansion of specific cellular subsets in older obese adults compared to young, including VA3 adipocytes, hASPC6 progenitors, and hMES1 mesothelial cells (black ovals). The endothelial cell subclusters were relatively enriched in middle-aged adults compared to young or old, suggesting an age-associated redistribution of the vascular compartment.

**Extended Figure 3**. **Senescent cells were most abundant in one to two subtypes of each cell-type in VAT of older obese adults. (a-o)** UMAP analysis to separate subtypes within each cell-type. Predicted SnCs indicated in red. Beneath each UMAP, SnCs are quantitated by cell subtype. SnCs, predicted by SEAC_UCELL, SEAC-REF. Cell Correlation or SenMayo, preferentially accumulated in cell-type subsets hASPC6/hASPC3, VA3/VA5, hENDOS2/5, hMES1, hMAC1/3 and monocytes1. Statistical significance was assessed using two-way ANOVA followed by Tukey's multiple-comparisons test, with age group and cellular subset as factors. The interaction term evaluates whether age-associated differences in predicted SnC accumulation vary among cellular subsets. The column factor (age group) evaluates the overall effect of age across cellular subsets. Statistical significance is indicated as $P < 0.05$ (*), $P < 0.01$ (**), $P < 0.001$ (***), and $P < 0.0001$ (****); ns, not significant.

**Extended Figure 4. Senescent cells occurred in the same cell subtypes in VAT of non-obese adults as in obese adults. (a-d)** UMAPs analysis to identify SnCs (red) in younger (18-35y) and older adults (52-75y) with low BMI (<30 kg/m²) (left panel). UMAP analysis to identify cell-type subtypes (middle panel). Expression of lineage markers in each of the cell-types used to discriminate subclusters of cells (right panel). Publicly available snRNA-seq datasets used for this analysis include those reported by Jung et al. (2026), Lazarescu et al. (2025), and Emont et al. (2022). The predicted SnCs were most abundant in hASPC6, hMES1, hENDO2, and hVA3 and all of which overlap with cellular subtypes with increase SnCs in obese individuals. Notably, SnCs were observed in the same cellular subtypes between young and old lean individuals, with greater abundance in the older subjects.

**Extended Figure 5. Age-associated signaling pathway remodeling in human VAT is most pronounced in older adults.** Examination of the concordance in the signaling pathways identified as altered between age groups of obese individuals. **(a)** GSEA analysis of bulkRNAseq data from older vs. young obese individuals. **(b)** Seurat analysis of snRNAseq data from older vs. young adult obese individuals. Pathways are indicated on the x-axis. Age-associated signaling molecule expression was concordant (75% similarity) between bulk- and snRNAseq datasets. **(c-d)** MSigDB Hallmark pathway enrichment analysis showed ~80% overlap in pathways altered between older versus younger and older versus middle-aged adults. **(e)** Hallmark pathway comparisons between middle-aged and younger adults showed no significant differences. **(f)** Comparison of $-\log_2$ P values across age group comparisons, showed similar pathway-level differences in older adults compared to middle-aged or younger adults.

**Extended Figure 6. Increased cell communication in VAT from older obese individuals.** CellChat analysis of snRNAseq data from young (YA), middle-aged (MA) and older obese adults (OA). **(a-c)** Heat maps indicate the strength of the incoming signals by pathway (y-axis) and cell-type (x-axis); colors are used to indicate distinct cell-types. Histogram above the heatmap illustrate the total predicted amount of incoming signal by cell-type. Shades of green are used to indicate the strength of the signal by cell-type. Grey bars to the right of the heatmap indicate the relative strength of that signal across all cell-types. Note the y-axes for OA is 2x greater than YA and MA. **(d-f)** The ligand-receptor signals related to inflammation predicted to by upregulated in older obese adults compared to young including CSF, MHC-1 and ICAM. The color of the lines indicates the cell source of the signal. The cells predicted to receive the paracrine and autocrine signals are indicated. The width of the line indicates the predicated strength of each signal.

**Extended Data 7. Senescent cells establish a conserved intercellular communication program.** CellChat Pattern Recognition Analysis of the snRNAseq data was used to identify recurring profiles of outgoing and incoming signals (patterns) by cell type using the SEAC-Reference cell correlation to define SnCs (**Figure 3a, b**). Here, the same analysis was repeated using SnCs identified by SenMayo **(a,b)** and SEAC-UCell **(c,d)**. Outgoing and incoming signaling interactions were each organized into six distinct communication patterns, conserved across all three methods of identifying SnCs but differing between the two signaling directions. The left most column identifies cell-types, non-senescent on top, senescent subclusters on the bottom (endo/endothelia; Mac/macrophage; Mes/mesothelial; VA/visceral adipocyte), which are predicted to be sending signals based on their transcriptomic profile **(a,c)** or receiving signals **(b,d)**. The second column indicates the pattern of signals emitted **(a,c)** or received **(b,d)**. The far-right column (grey boxes) indicates the genes defining each pattern.

**Extended Figure 8. Senescence-associated WNT signaling and KDM6B expression implicated in altered adipogenic regulation of aging human visceral adipose tissue. (a-c)** CellChat ligand-receptor analysis of snRNAseq data predicted SnCs identified using three complementary approaches: SenMayo, SEAC-reference cell correlation, and SEAC-UCell-revealed WNT9A-LRP6 as a prominent predicted signaling interaction across multiple SnC lineages in older obese adult VAT. **(d-f)** qPCR measure of *TNF*, $p21^{CIP1}$ and $p16^{INK4a}$, respectively, in older obese adult VAT relative to the average expression in younger obese adults. Each dot represents a biological replicate. One-way ANOVA with post-hoc Tukey's multiple-comparisons tests; p values for OA vs. YA are indicated above the graph. **(g-j)** Increased expression of *KDM6B* and *WNT9A* was detected in the cell-type subclusters identified as enriched in SnCs in older obese adults. *PPARG* was downregulated across adipocyte populations suggesting altered adipogenic regulation. **(k-m)** Spatial protein expression of the endothelial marker CD31, HIF1A, γH2AX, and CD38, assessed using the COMET spatial proteomics platform. **(n-s)** Visium HD data analysis reveals a trend toward increased expression of *KDM6B* and *p21/CDKN1A*, and across cell-types in VAT from older adults. In contrast, *p16/CDKN2A* expression was increased only in adipocytes and whole VAT, but not in the other cell-types. **(p,q)** Representative Loupe Browser images from Visium HD spatial transcriptomics showed the spatial distribution of *KDM6B* expression (blue) in mesothelial and endothelial cells within VAT.
