## Extended Figures for "Aging drives senescence in visceral adipose to a greater extent than obesity, instigating multicellular common signaling patterns that impair organ function and drive inflammation"

**Extended Figure 1**

Inflammation

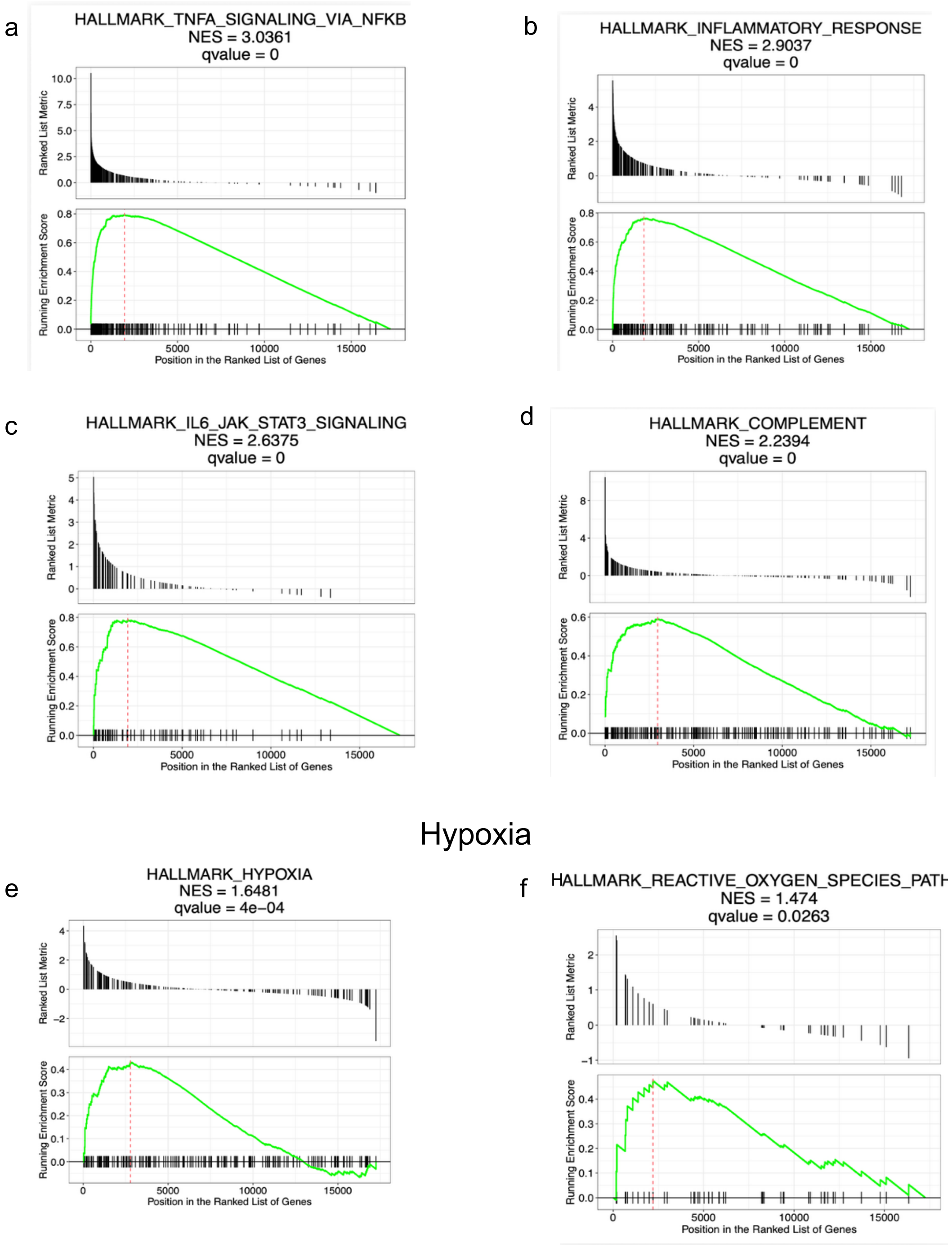

**Continued Extended Figure 1**

#### DNA damage

g
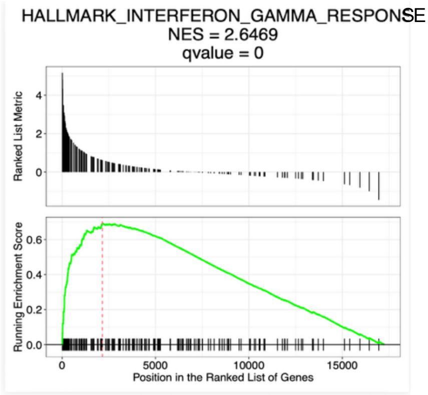


h
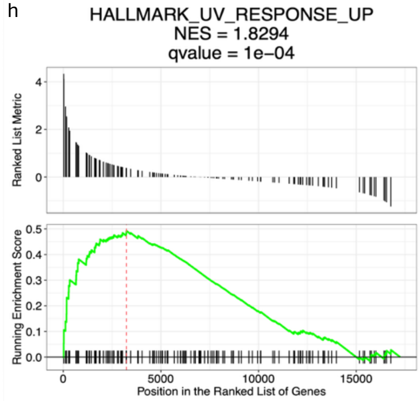


i
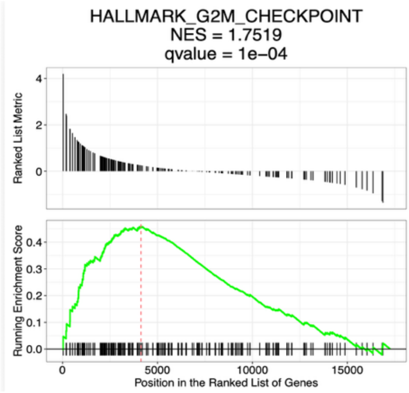


j
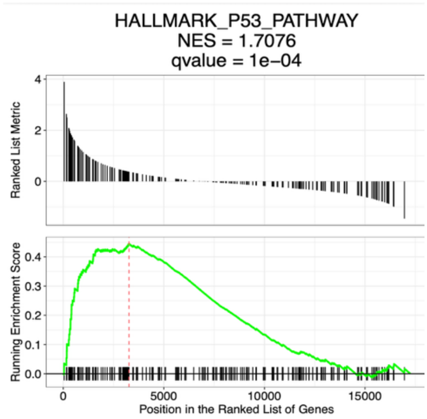


#### Fibrosis

k
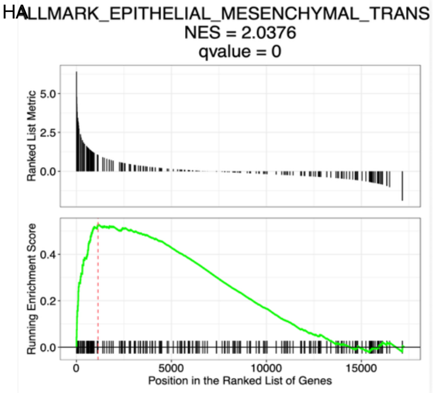


l
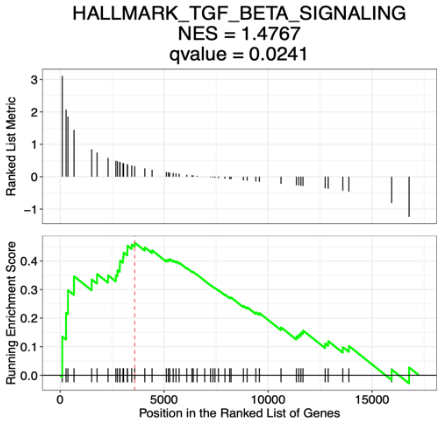

### Extended Figure 2

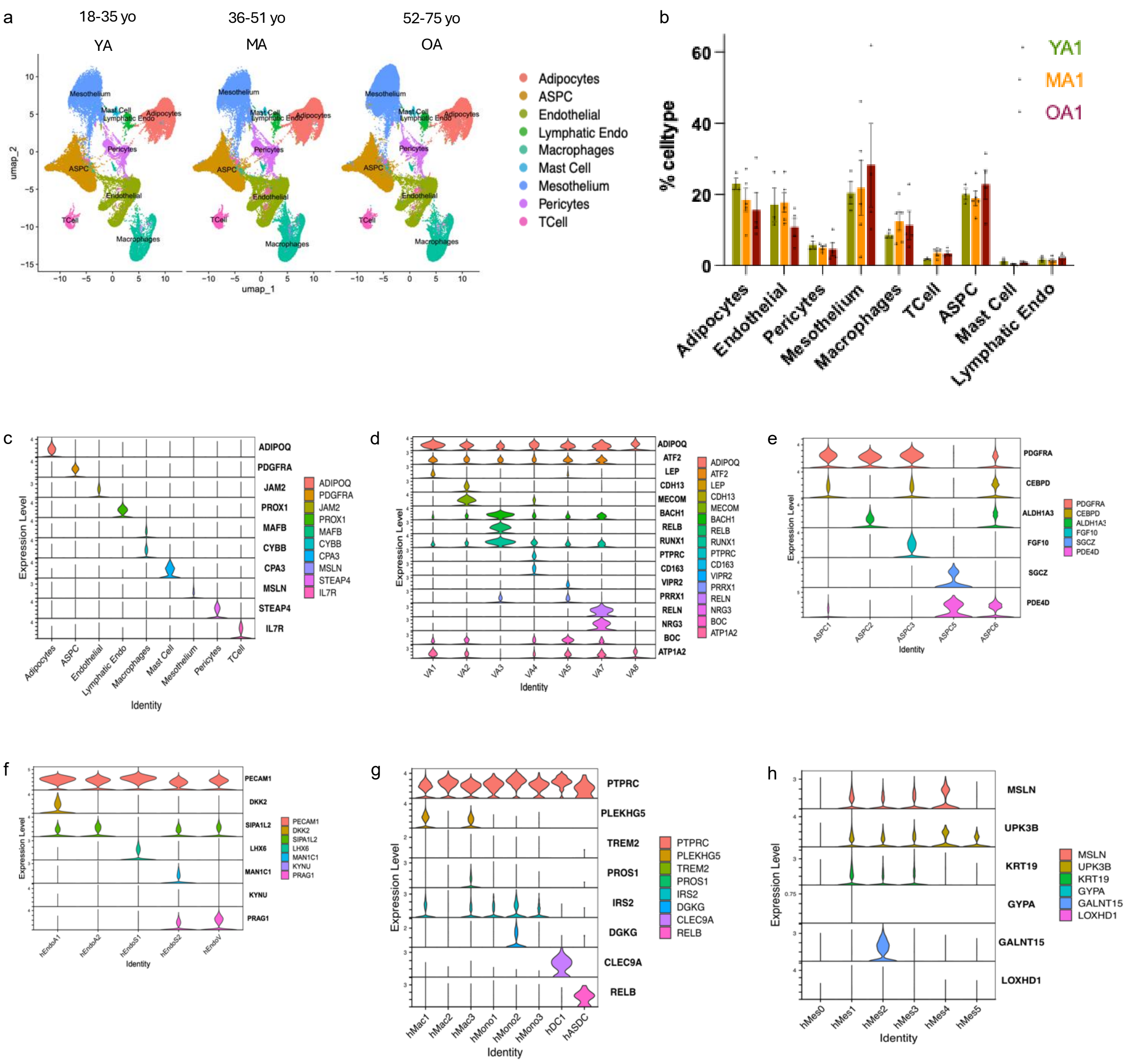

**Continued Extended Figure 2**

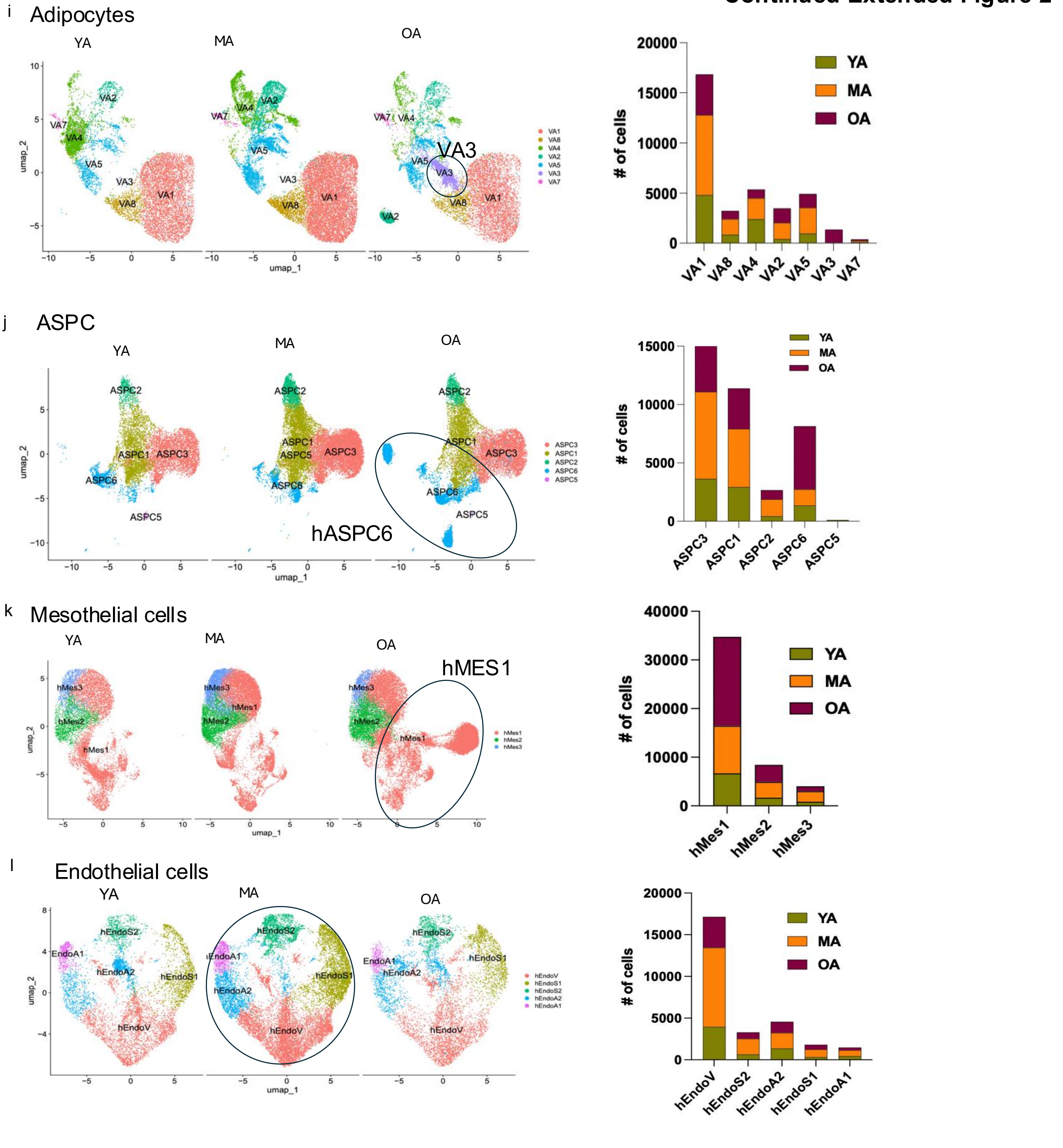

#### Extended Figure 3

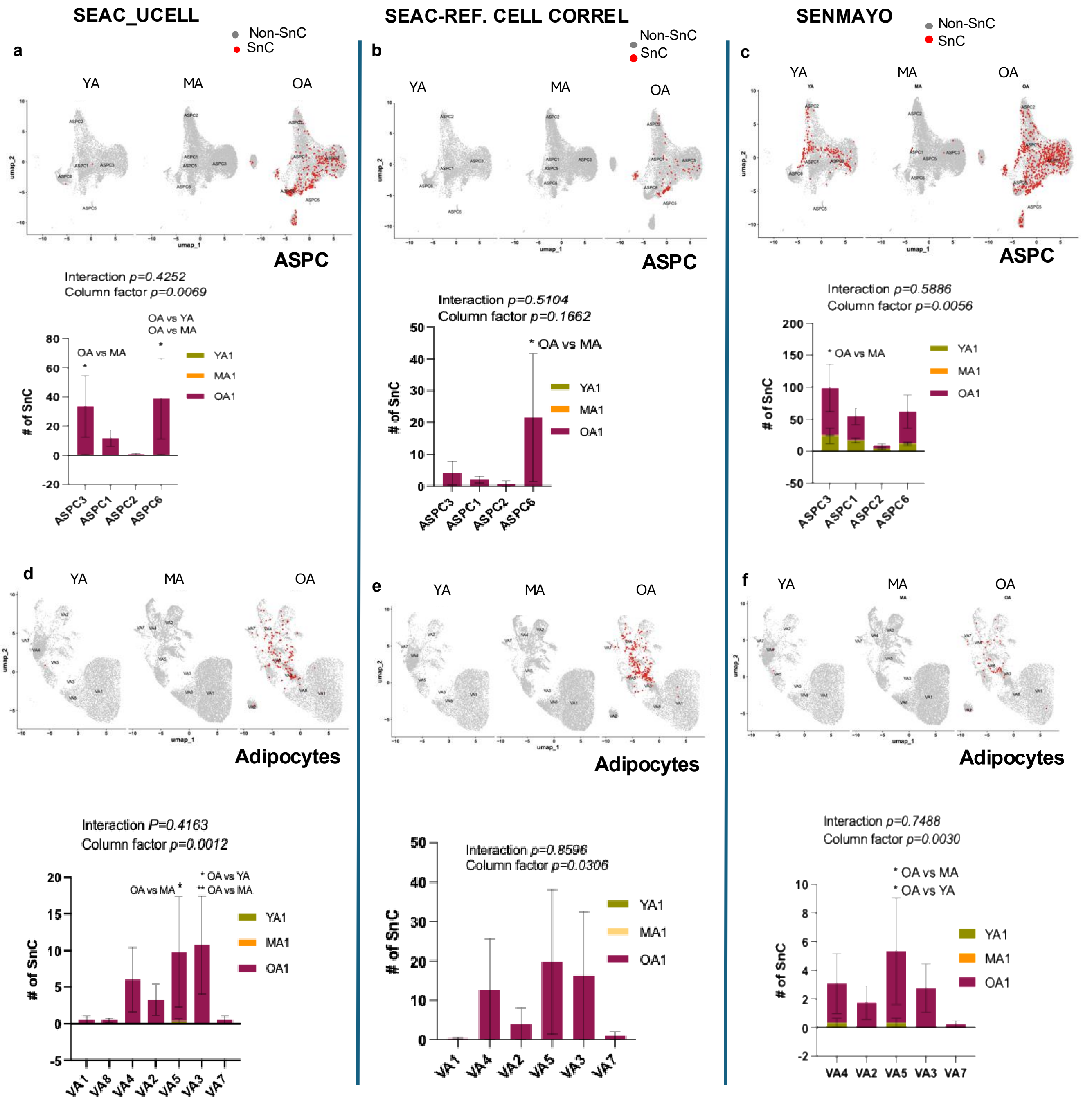

**Continued Extended Figure 3**

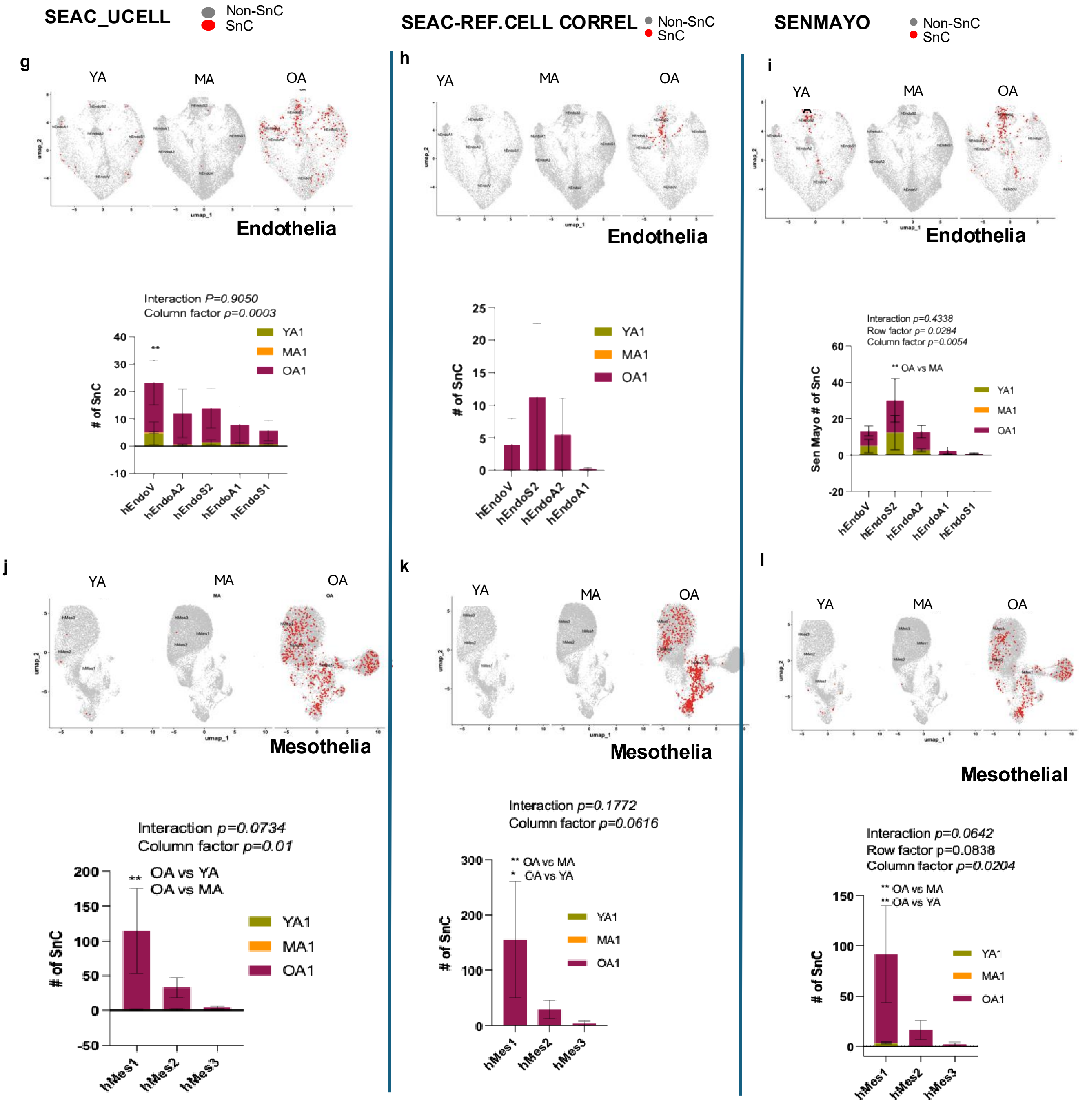

### Continued Extended Figure 3

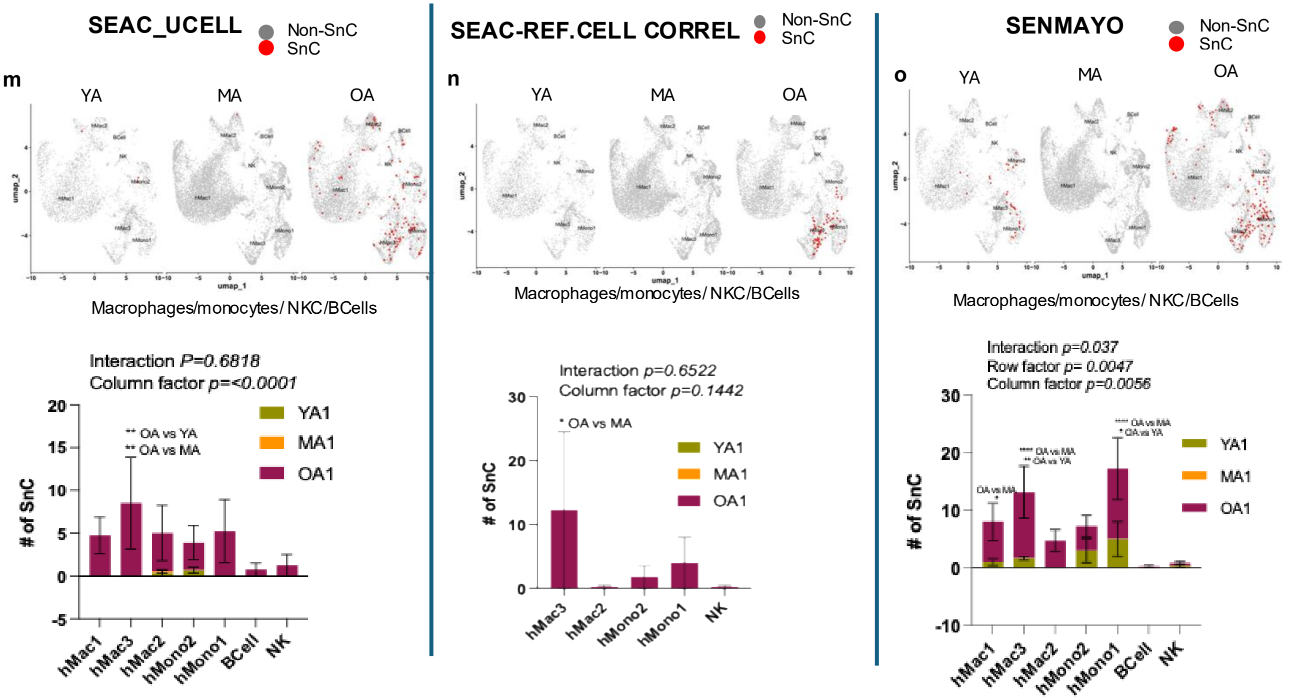

#### Extended Figure 4

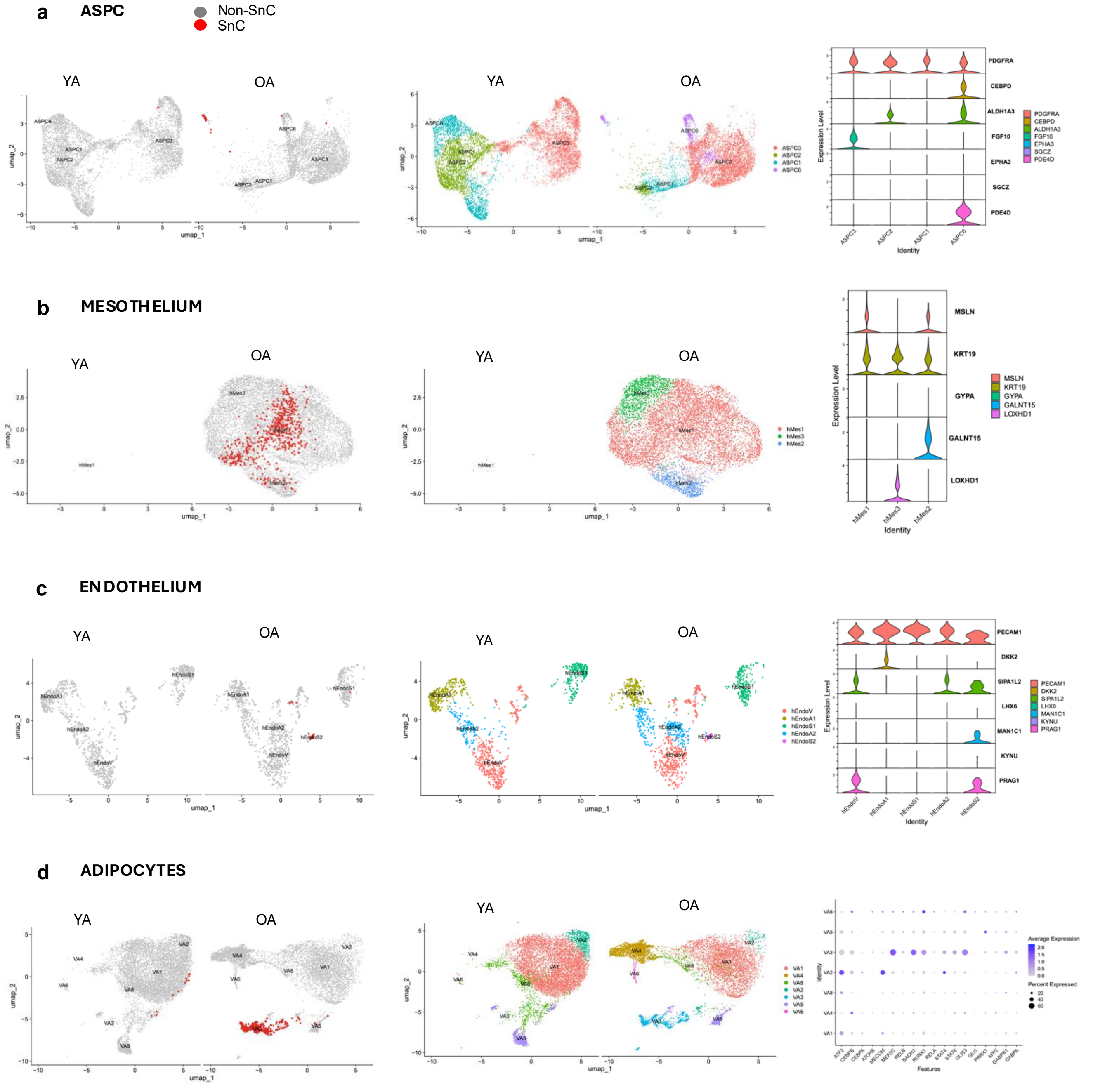

**Extended Figure 5**

**a** OA vs YA: Bulk RNAseq

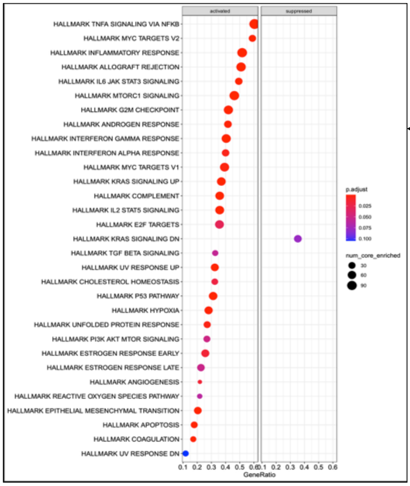


**b** OA vs YA: sn-RNAseq

75%

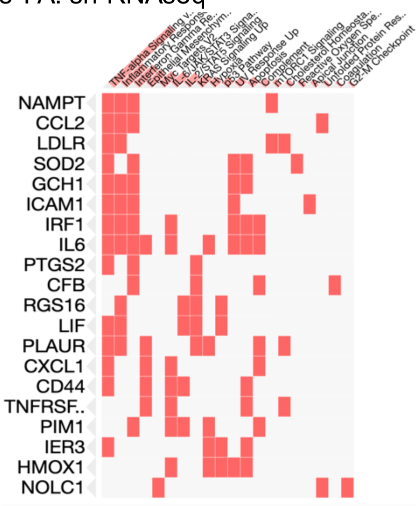


**c** OA vs YA - snRNAseq

| Index | Name | P-value | Adjusted p-value | Odds Ratio | Combined score |
|---|---|---|---|---|---|
| 1 | TNF-alpha Signaling via NF-kB | 1.609e-33 | 6.596e-32 | 25.24 | 1906.05 |
| 2 | Inflammatory Response | 1.755e-12 | 3.598e-11 | 10.66 | 288.57 |
| 3 | Interferon Gamma Response | 1.863e-11 | 2.546e-10 | 9.96 | 246.03 |
| 4 | Epithelial Mesenchymal Transition | 1.467e-8 | 1.504e-7 | 7.94 | 143.17 |
| 5 | Myc Targets V2 | 1.086e-7 | 8.902e-7 | 16.46 | 263.92 |
| 6 | IL-6/JAK/STAT3 Signaling | 2.185e-7 | 0.000001493 | 11.91 | 182.72 |
| 7 | IL-2/STAT5 Signaling | 8.067e-7 | 0.000004725 | 6.69 | 93.93 |
| 8 | KRAS Signaling Up | 0.000005689 | 0.00002916 | 6.04 | 72.93 |
| 9 | Hypoxia | 0.0001900 | 0.0007251 | 4.84 | 41.45 |
| 10 | p53 Pathway | 0.0001900 | 0.0007251 | 4.84 | 41.45 |
| 11 | UV Response Up | 0.0001945 | 0.0007251 | 5.46 | 46.64 |
| 12 | Apoptosis | 0.0002212 | 0.0007558 | 5.35 | 45.03 |
| 13 | Complement | 0.0009341 | 0.002946 | 4.26 | 29.68 |
| 14 | mTORC1 Signaling | 0.004067 | 0.01191 | 3.68 | 20.28 |
| 15 | Cholesterol Homeostasis | 0.006473 | 0.01769 | 5.75 | 28.99 |
| 16 | Reactive Oxygen Species Pathway | 0.01295 | 0.03319 | 6.54 | 28.42 |
| 17 | Apical Junction | 0.01549 | 0.03737 | 3.13 | 13.03 |

80%

**d** OA vs MA - snRNAseq

| Index | Name | P-value | Adjusted p-value | Odds Ratio | Combined score |
|---|---|---|---|---|---|
| 1 | TNF-alpha Signaling via NF-kB | 1.116e-43 | 4.685e-42 | 32.84 | 3247.60 |
| 2 | Epithelial Mesenchymal Transition | 1.697e-13 | 2.376e-12 | 11.32 | 332.72 |
| 3 | Inflammatory Response | 1.697e-13 | 2.376e-12 | 11.32 | 332.72 |
| 4 | Interferon Gamma Response | 1.991e-10 | 2.090e-9 | 9.22 | 205.95 |
| 5 | IL-6/JAK/STAT3 Signaling | 1.155e-9 | 9.701e-9 | 15.02 | 309.19 |
| 6 | IL-2/STAT5 Signaling | 8.506e-7 | 0.000005382 | 6.66 | 93.07 |
| 7 | Hypoxia | 8.970e-7 | 0.000005382 | 6.62 | 92.22 |
| 8 | Myc Targets V2 | 0.000001817 | 0.000009540 | 13.97 | 184.68 |
| 9 | Apoptosis | 0.000005467 | 0.00002506 | 6.81 | 82.55 |
| 10 | Complement | 0.000005966 | 0.00002506 | 6.01 | 72.26 |
| 11 | UV Response Up | 0.00003229 | 0.0001233 | 6.18 | 63.93 |
| 12 | KRAS Signaling Up | 0.00003610 | 0.0001264 | 5.40 | 55.27 |
| 13 | Coagulation | 0.00007872 | 0.0002543 | 6.27 | 59.26 |
| 14 | p53 Pathway | 0.0001972 | 0.0005917 | 4.81 | 41.05 |
| 15 | Allograft Rejection | 0.0009646 | 0.002701 | 4.23 | 29.39 |
| 16 | Myc Targets V1 | 0.004179 | 0.01097 | 3.67 | 20.08 |
| 17 | Angiogenesis | 0.005594 | 0.01331 | 9.08 | 47.07 |
| 18 | Unfolded Protein Response | 0.005703 | 0.01331 | 4.65 | 24.03 |
| 19 | Cholesterol Homeostasis | 0.006587 | 0.01456 | 5.72 | 28.74 |
| 20 | Estrogen Response Early | 0.01584 | 0.03024 | 3.11 | 12.89 |
| 21 | mTORC1 Signaling | 0.01584 | 0.03024 | 3.11 | 12.89 |
| 22 | Xenobiotic Metabolism | 0.01584 | 0.03024 | 3.11 | 12.89 |
| 23 | TGF-beta Signaling | 0.01704 | 0.03111 | 5.87 | 23.89 |

**e** MA vs YA - snRNAseq

| Index | Name | P-value | Adjusted p-value | Odds Ratio | Combined score |
|---|---|---|---|---|---|
| 1 | Complement | 0.06031 | 0.7730 | 2.81 | 7.88 |
| 2 | Xenobiotic Metabolism | 0.4335 | 0.7730 | 1.37 | 1.14 |
| 3 | Allograft Rejection | 0.4335 | 0.7730 | 1.37 | 1.14 |
| 4 | KRAS Signaling Up | 0.4335 | 0.7730 | 1.37 | 1.14 |
| 5 | KRAS Signaling Dn | 0.4335 | 0.7730 | 1.37 | 1.14 |
| 6 | Interferon Gamma Response | 0.4335 | 0.7730 | 1.37 | 1.14 |
| 7 | Coagulation | 0.6400 | 0.7730 | 0.99 | 0.44 |
| 8 | UV Response Up | 0.6897 | 0.7730 | 0.86 | 0.32 |
| 9 | Mitotic Spindle | 0.7713 | 0.7730 | 0.68 | 0.18 |
| 10 | IL-2/STAT5 Signaling | 0.7713 | 0.7730 | 0.68 | 0.18 |

**f**

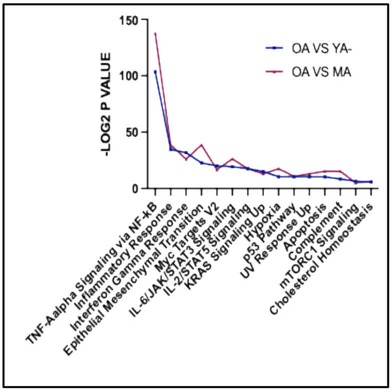

**Extended Figure 6**

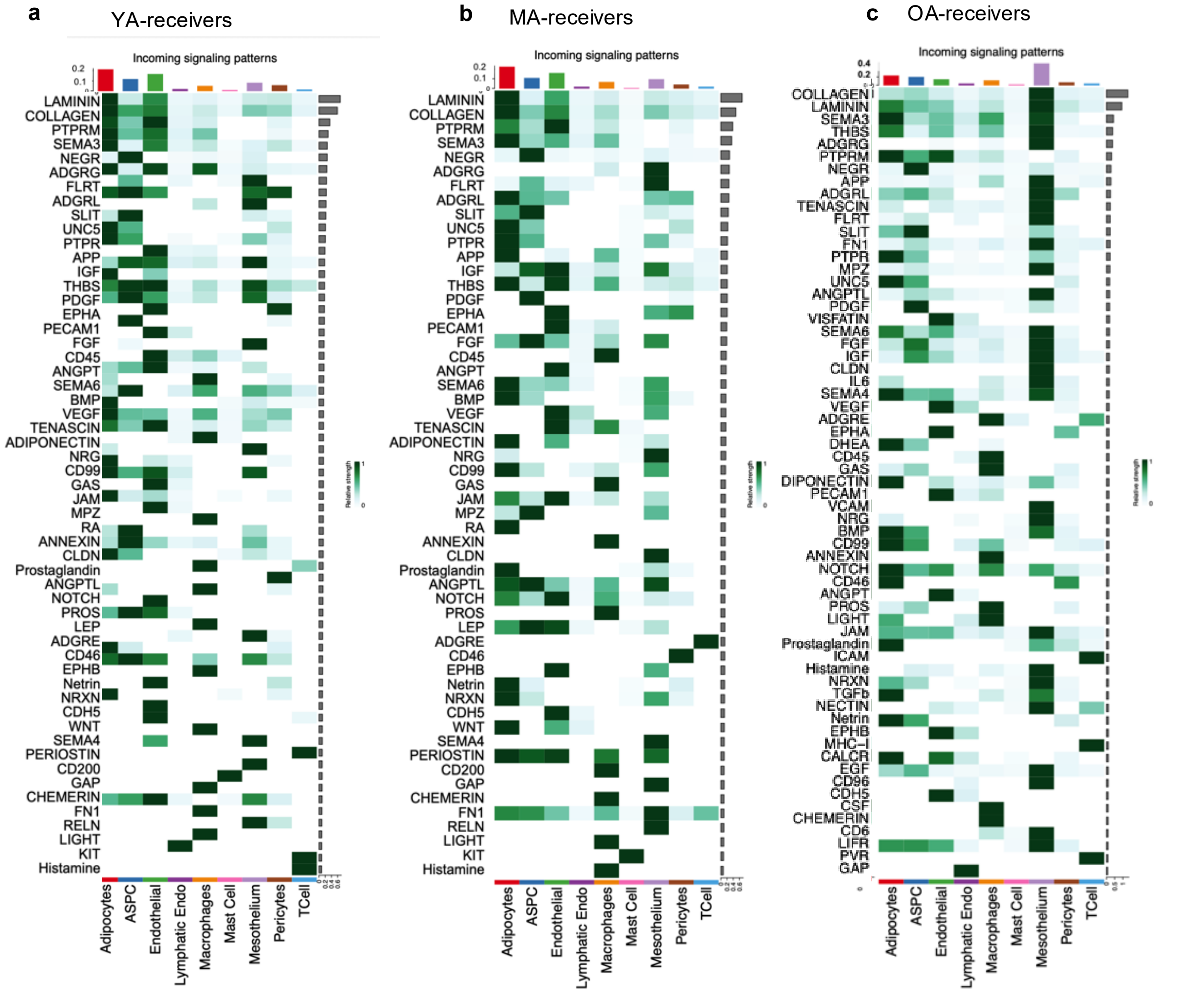

**Continued Extended Figure 6**

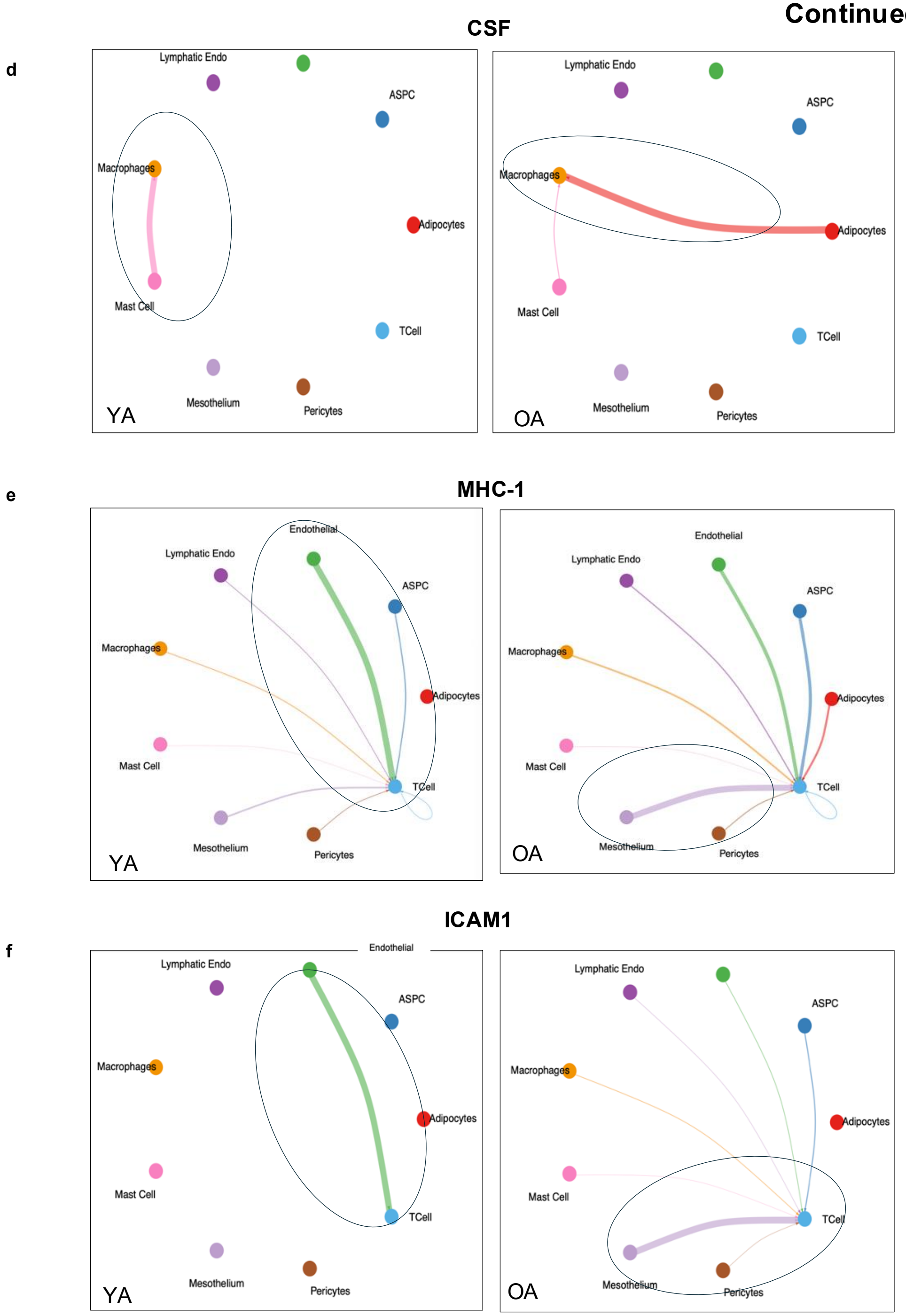

### Extended Figure 7

#### SenMayo gene panel

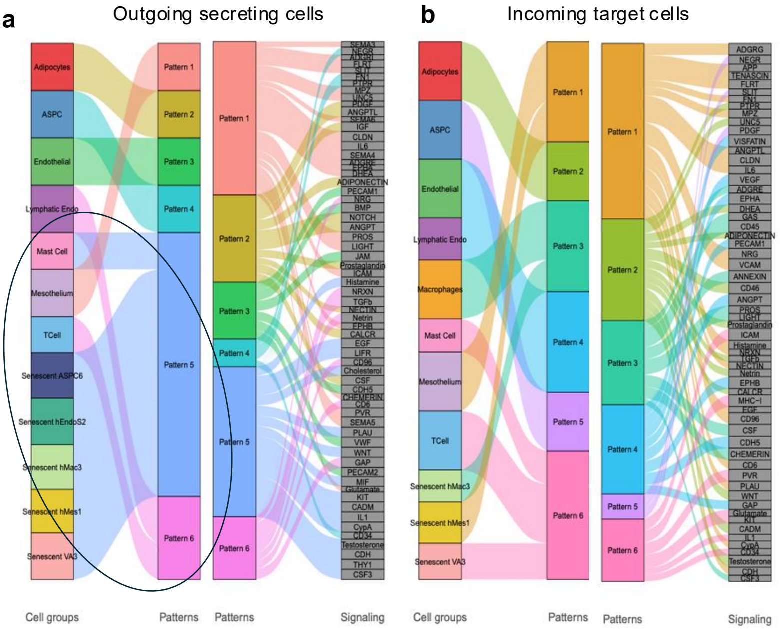


#### SEAC-UCELL

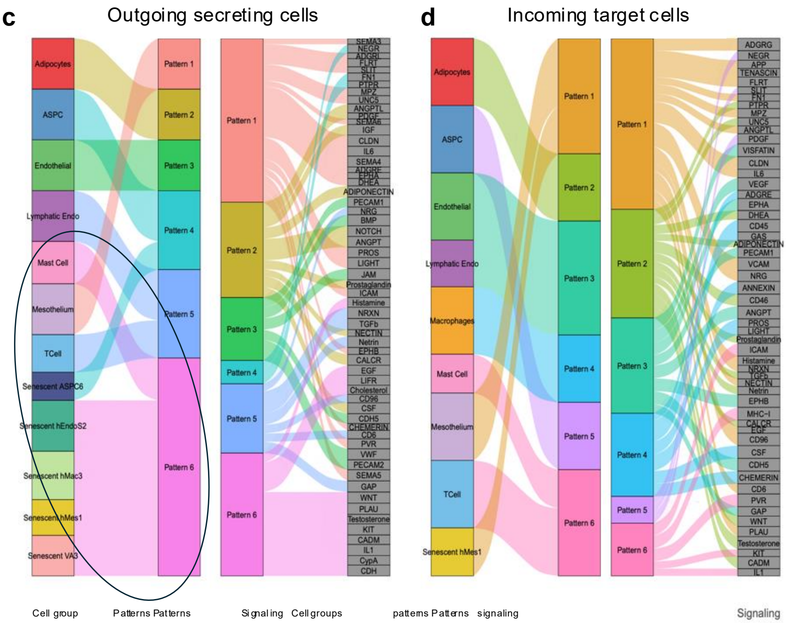

### Extended Figure 8

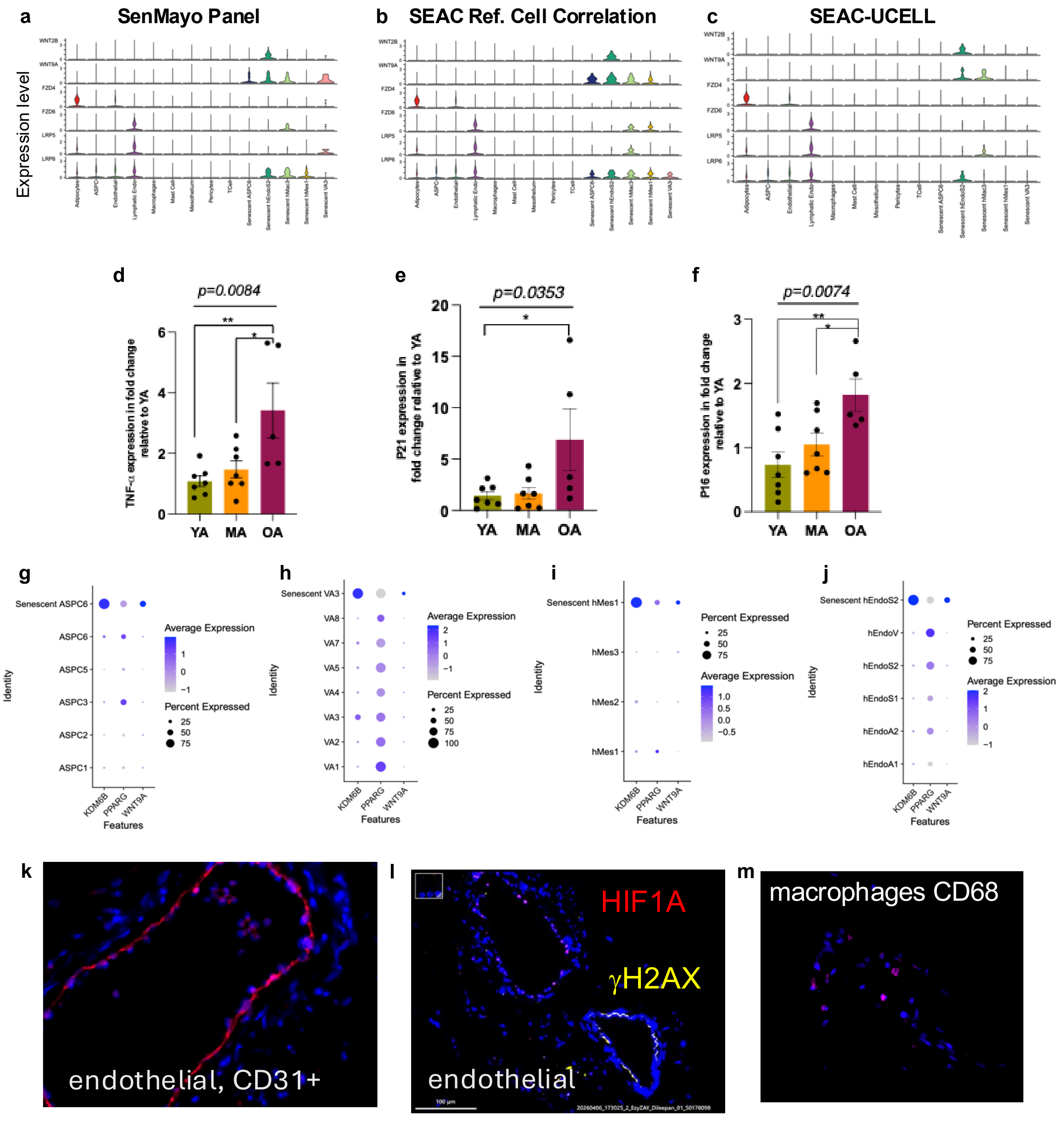

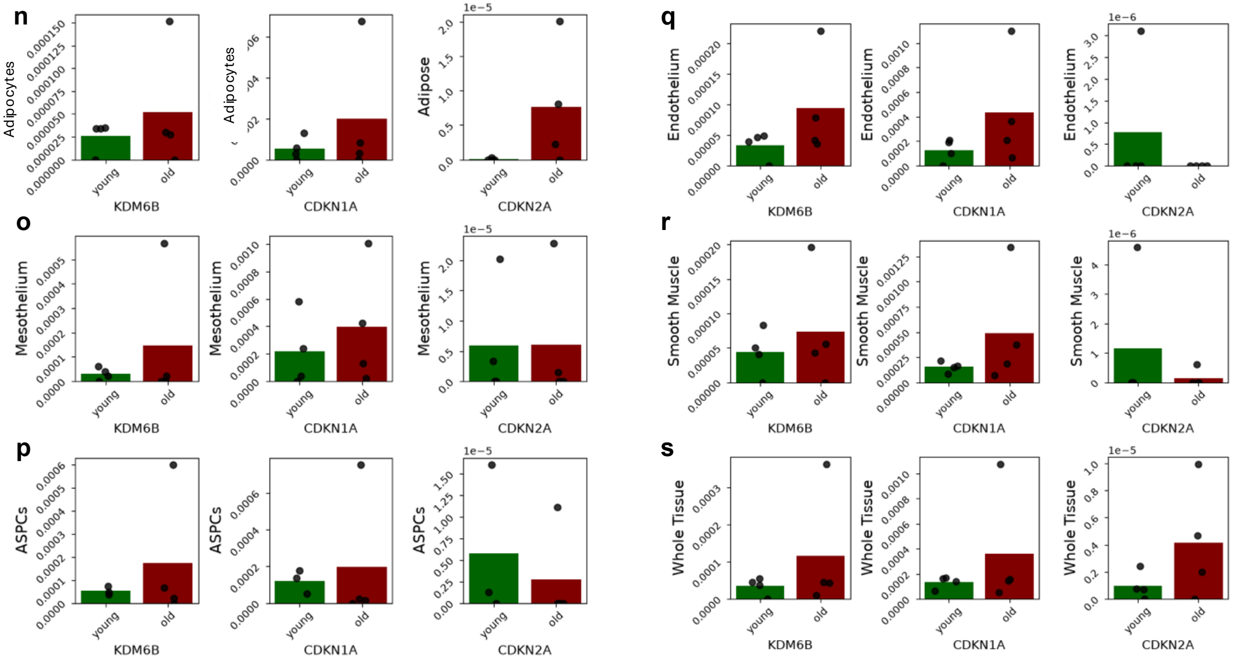
n
Adipocytes
Adipose
o
Mesothelium
p
ASPCs
q
Endothelium
r
Smooth Muscle
s
Whole Tissue
young
old
KDM6B
CDKN1A
CDKN2A
1e−5
1e−6

p

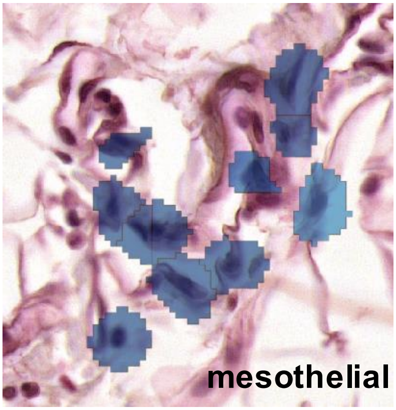
mesothelial

q

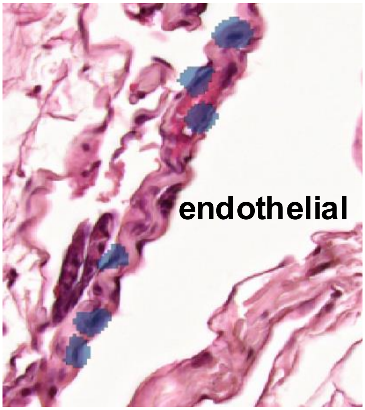
endothelial
